# Genetic factors driving multi-host infection in a core member of the root mycobiota

**DOI:** 10.1101/2025.12.01.690973

**Authors:** Ram-Sevak Raja-Kumar, Fantin Mesny, Arpan Kumar Basak, Jacy Newfeld, Guillaume Chesneau, Frederickson Entila, Tak Lee, Linda Rigerte, Stephanie Carvajal Acevedo, Bruno Hüttel, Pedro W. Crous, Jose G. Maciá-Vicente, Helen Stewart, Matthew Ryan, Ahmad M. Fakhoury, Soledad Sacristán, Isabelle Batisson, Stefano Dumontet, Wade H. Elmer, Jana Henzelyová, Joanna S. Kruszewska, Jessica M. Nelson, Cara M. Santelli, Markus Pauly, Antonio Molina, Kei Hiruma, Stéphane Hacquard

**Affiliations:** Max Planck Institute for Plant Breeding Research, 50829 Cologne, Germany; Malopolska Centre of Biotechnology, Jagiellonian University, Krakow, 30-387 Poland; Department of Life Sciences, Multidisciplinary Sciences, Graduate School of Arts and Sciences, The University of Tokyo, 3-8-1, Komaba, Meguro-ku, Tokyo 153-8902, Japan; Max Planck Genome Centre Cologne, Max Planck Institute for Plant Breeding Research, 50829 Cologne, Germany; Westerdijk Fungal Biodiversity Institute, P.O. Box 85167, 3508 AD Utrecht, The Netherlands; Microbiology, Department of Biology, Faculty of Science, Utrecht University, Padualaan 8, 3584 CT Utrecht, The Netherlands; Department of Genetics, Biochemistry and Microbiology, Forestry and Agricultural Biotechnology Institute (FABI), University of Pretoria, Pretoria, 0002, South Africa; Marine Sciences and Applied Biology, University of Alicante, PO Box 99, 03083 Alicante, Spain; CABI, Imperial site at Silwood Park, Buckhurst Rd, Ascot SL5 7PY, United Kingdom; School of Agricultural Sciences, Southern Illinois University, Carbondale, IL 62901, USA; Centro de Biotecnología y Genómica de Plantas, Universidad Politécnica de Madrid (UPM)-Instituto Nacional de Investigación y Tecnología Agraria y Alimentaria (INIA/CSIC), Campus Montegancedo-UPM, 28223-Pozuelo de Alarcón (Madrid), Spain; Departamento de Biotecnología-Biología Vegetal, Escuela Técnica Superior de Ingeniería Agronómica, Alimentaria y de Biosistemas, Universidad Politécnica de Madrid, 28040-Madrid, Spain; Université Clermont-Auvergne, CNRS, Laboratoire Microorganismes: Génome et Environnement, F-63000 Clermont-Ferrand, France; Department of Sciences and Technologies, University of Napoli ‘Parthenope’, Centro Direzionale, Isola C4, 80143 Napoli, Italy; Department of Plant Pathology and Ecology, The Connecticut Agricultural Experiment Station, New Haven, Connecticut 06504, USA; Department of Genetics, Institute of Biology and Ecology, Faculty of Science, Pavol Jozef Šafárik University in Košice, Mánesova 23, 04154, Košice, Slovakia; Institute of Biochemistry and Biophysics, Polish Academy of Sciences, Warsaw, Poland; Natural History Museum, University of Oslo, 0562 Oslo, Norway; Department of Earth and Environmental Sciences, University of Minnesota Twin Cities, Minneapolis, MN, USA; Institute of Plant Cell Biology and Biotechnology, Heinrich Heine University Düsseldorf, Düsseldorf, Germany; Cluster of Excellence on Plant Sciences (CEPLAS), Max Planck Institute for Plant Breeding Research, 50829 Cologne, Germany

## Abstract

Core members of the fungal root microbiota include pathogens capable of colonizing multiple hosts, yet the underlying genetic determinants remain unknown. We report that *Plectosphaerella cucumerina* is a core member of the *Arabidopsis thaliana* root microbiota displaying high pathogenic potential and multi-host colonization capabilities. Establishment of a *Plectosphaerella* reference culture collection, followed by whole-genome sequencing of 72 strains reveals subtle phenotypic and genotypic variation that associate with fungal phylogeny, but not host plant identity. Transcriptome profiling of a model *P. cucumerina* isolate in roots of multiple hosts identifies core and host-specific fungal processes linked to carbon catabolism and root cell wall deconstruction of the hosts. A fungal gene encoding a candidate β-1,3-glucanase (GH64) was identified as a key genetic factor driving infection and disease in plants that diverged 110 million years ago. The gene is enriched in plant-colonizing fungi and consistently functions as a disease determinant in the root pathogen *Colletotrichum incanum*. We conclude that diverse and tunable fungal repertoires of carbohydrate-active enzymes act as disease determinants and drive multi-host compatibility belowground.

## Introduction

The root mycobiota – the community of fungi that associate with the roots of plants – include members that establish beneficial relationships but also isolates that conditionally cause diseases or hamper plant performance^1, 2, 3, 4, 5, 6^. Consistently, several core fungal taxa that reproducibly colonize roots of healthy plants in natural populations are often harmful under laboratory conditions^7, 8, 9^, illustrating that the pathogenic lifestyle of several root mycobiota members is often kept at bay in nature. In *Arabidopsis thaliana*, it has been reported that the protective activity of bacterial root commensals is at least as important as the innate immune branch involving tryptophan-derived specialized metabolites for controlling fungi in roots^4, 9, 10, 11^, thereby maintaining microbial interkingdom balance for plant health^12^.

Given that soil and root-associated fungal communities are extensively shaped by the local climatic, edaphic and host conditions^13, 14, 15^, high endemism in fungal communities has been repeatedly reported^16, 17^, with very few core fungal species detected worldwide^18^. Widespread fungal root colonizers appear particularly successful due to their ability to efficiently disperse, infect multiple hosts, and plastically adapt to various environmental conditions^19^. However, the underlying molecular mechanisms enabling adaptation to a broad range of hosts and environments in root mycobiota members remains unknown.

Plant-associated fungi produce remarkable cocktails of carbohydrate-active enzymes (CAZymes) that target plant and/or microbial cell walls and outer layers. These enzymes facilitate fungal entry into plant tissues, promote carbohydrate extraction from host cell walls or modulate immune responses^8, 20, 21, 22, 23^, but can also contribute to antagonistic microbe-microbe interactions^24, 25^. The repertoire of fungal CAZymes often explains fungal lifestyles or correlates with adaptation to specific hosts or organs^26, 27, 28, 29, 30, 31^. For example, transitions from saprotrophic fungi to ectomycorrhizal symbionts have occurred multiple times independently, involving significant losses of CAZymes involved in lignin and cellulose deconstruction^32, 33^. In contrast, endophytism in Arabidopsis root mycobiota members does not correlate with a reduction of saprotrophic traits^3, 8, 11, 34^ but involves pectin-degrading enzymes that modulate fungal root colonization aggressiveness^8^. However, the extent to which the diversity, composition, and transcriptional activation of CAZyme repertoires contribute to fungal adaptation to multiple hosts and environments in root mycobiomes remains unclear.

We report that the fungus *Plectosphaerella cucumerina*^35^ – a well-known pathogen of diverse plant species causing wilt disease, root rot disease or leaf necrosis^36, 37, 38, 39^ – is also a prevalent species colonizing the roots of *A. thaliana* at a continental scale. The fungus (phylum Ascomycota, class Sordariomycetes, order Glomerellales, family Plectosphaerellaceae) exhibits broad host range colonization capabilities and acts as a root pathogen of the dicotyledonous plants *Solanum lycopersicum* (tomato) and *A. thaliana* but not of the monocotyledonous host *Hordeum vulgare* (barley). We assembled a large collection of 69 *P. cucumerina* isolates that were retrieved from various host plants worldwide as well as high quality – long read-based – genome assemblies. Multi-host recolonization experiments using the model *P. cucumerina* 0016 strain, followed by fungal transcriptome sequencing revealed 1) high plasticity in catabolic process activation that mirrors root cell wall composition of the cognate hosts, 2) the key contribution of a *P. cucumerina* β-1,3-glucanase (glycoside hydrolase familly 64, GH64) for driving multi-host infection and disease in host plants that diverged 110 million years ago (MyA) and 3) the evolutionary conservation of GH64 as a root infection determinant in the root pathogen *Colletotrichum incanum* (order Glomerellales, family Glomerellaceae). Our results indicate that adaptation to multiple hosts across various environments is conferred by a combination of genome-rich CAZyme repertoires and the plastic up-regulation of these genes in response to both conserved and host-specific cell wall cues.

## Results

### *P. cucumerina* is a core member of the *A. thaliana* root mycobiota and has a broad host range

We hypothesized that specific root mycobiota members can reproducibly colonize roots of healthy Arabidopsis in nature, irrespective of differences in climate and soil conditions. To test for the presence of a core root mycobiota, we first used amplicon sequencing data from two previously published studies^9, 14^ (18 sites, n = 291 root samples in total) and re-analyzed them at amplicon sequence variant (ASV)-level resolution (i.e., Zero-radius operational taxonomic unit^40^, Zotu, see methods). From the 26,976,402 sequenced ITS reads, we identified 338 Zotus that were detected in *A. thaliana* roots in at least one of the 18 sites across Europe [mean relative abundance (RA) > 0.1%, **Fig. 1a** and **Supplementary Table 1**]. Notably we identified 5 Zotus that were reproducibly detected in more than 80% of the 291 root samples, thereby constituting a small – yet multispecies – core Arabidopsis root mycobiota (**Fig. 1a**). These corresponded to Zotu2 (*Fusarium* sp.), Zotu9 (*Plectosphaerella* sp.), Zotu1 (*Dactylonectria* sp.), Zotu6 (*Fusarium* sp.) and Zotu3 (*Fusarium* sp.) that have occurrence values of 1, 0.98, 0.92, 0.87 and 0.87, respectively (**Fig. 1a** and **Supplementary Table 1**). Cross referencing with 488 ITS sequences corresponding to cultured isolates established from four of these sites (see methods, **Supplementary Table 1**) confirmed that multiple (>10) representative isolates had 100% sequence match to these Zotus (**Supplementary Fig. 1a**). Particularly, 106 ITS sequences from cultured isolates matched Zotu9 and were unambiguously assigned to *Plectosphaerella cucumerina* (Phylum: Ascomycota; class: Sordariomycetes, family: Plectosphaerellaceae; based on the RDP classifier^41^, **Supplementary Table 2, Supplementary Fig. 1b**), a well-known pathogen of several plant species, including Arabidopsis^37, 42^. We then used *P. cucumerina* as an input species in GlobalFungi^43^ and detected this species across a variety of niches (**Fig. 1b**) and particularly in soil, mosses, topsoil, rhizosphere and root/rhizosphere samples (pairwise one-sided Fisher’s exact tests, Benjamini-Hochberg correction, **Fig. 1c**), thereby confirming its presence at the soil-root interface. Root and/or rhizosphere samples in which *P. cucumerina* was detected were distributed worldwide across latitudes/longitudes and ecosystems (**Fig. 1d, Supplementary Fig. 2a,b**) and inspection of the corresponding plants revealed broad associations with a remarkable diversity of hosts, including monocots and dicots (**Fig. 1e**). The results indicate that *P. cucumerina* is a widespread root colonizer that has a broad range of hosts.

**Fig. 1:**
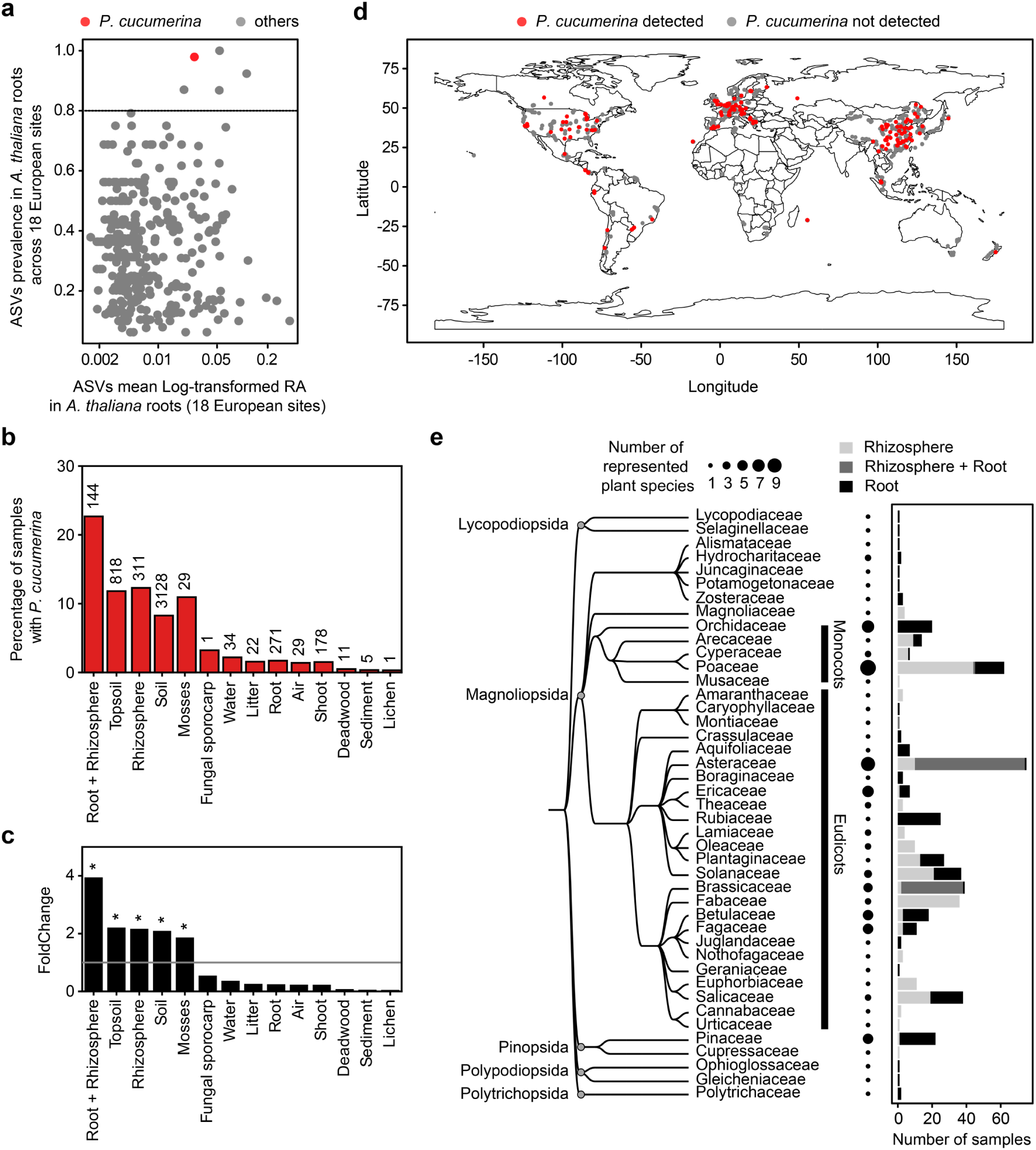
*P. cucumerina* is a core member of the *A. thaliana* root mycobiota and a widespread multi-host root colonizer. **a**, Re-analysis of amplicon sequence variants (i.e., zero radius OTUs, Zotus) sequenced from *A. thaliana* root samples (n = 281) from two previous studies^9, 14^. Zotus are plotted according to their prevalence and mean log-transformed relative abundance (RA) in roots across all samples. RA was calculated across samples where Zotus were detected (minimum 0.1% per sample). Only samples having more than 1,000 reads were considered for the analysis. The five Zotus above the dashed line corresponds to the most widespread root-colonizing taxa. *P. cucumerina* zOTU is highlighted in red. **b**, Analysis of GlobalFungi data^43^ showing detection of *P. cucumerina* in a variety of niches (n = 4,982/84,844 samples). Percentage of samples in which the *P. cucumerina* species was detected are plotted for each niche, with the corresponding numbers of samples annotated on the barplot. **c**, Fold-change values corresponding to the proportion of samples from a given niche in which *P. cucumerina* was detected divided by the proportion of samples in all other niches in which *P. cucumerina* was detected. Asterisks highlight the niches in which detection of *P. cucumerina* is significantly overrepresented (pairwise one-sided Fisher’s exact tests + Benjamini-Hochberg correction) **d**, Geographic origin of root and rhizosphere samples of the GlobalFungi database (n = 19,030 samples), with samples in which species *P. cucumerina* was detected in red (n = 726 samples). **e**, Phylogeny (family level) of the plant hosts associated to the GlobalFungi root and rhizosphere samples in which *P. cucumerina* was detected (n = 726 samples). The phylogenetic tree was assembled based on species names and NCBI Taxonomy data using Taxallnomy^82^. A bubble plot shows the number of species represented in each family. The stacked barplot highlights the number of samples associated to this plant host family in which *P. cucumerina* was detected, distinguishing root, rhizosphere and root + rhizosphere samples.

### Genetic and phenotypic differentiation among *P. cucumerina* isolates is explained by phylogeny, but not by the host of isolation

We hypothesized that high genomic and phenotypic variation explains broad fungal distribution and host range. We assembled a collection of 72 *Plectosphaerella* strains that were isolated from diverse plant families (including bryophytes^44^) and environments worldwide. Among these 69 belong to the *P. cucumerina* species complex, and three belong to the closely related species *P. plurivora* (n = 2) and *P. delsorboi* (n = 1) (**Supplementary Table 3**, **Fig. 2a**). PacBio long-read genome sequencing resulted in high-quality assemblies (sizes: 35.5 − 40.5 Mbp, median = 36.4 Mbp; number of contigs: 9 – 188, median = 27.5; N50: 0.77 – 7.66 Mbp, median = 2.69 Mbp) that contained similar sets of predicted genes (9,992 – 12,956 genes, median: 11,581.5) (**Supplementary Fig. 3a-d**). Phylogeny reconstruction using 5,466 single copy orthologs identified two *P. cucumerina* subspecies (subsp. 1: 39 strains, subsp. 2: 26 strains) and a *P. cucumerina* outgroup consisting of four strains (**Fig. 2a**, **Supplementary Fig. 4a-d**). A burst in transposon and retrotransposon elements was observed in subsp. 2 (**Supplementary Fig. 5a-e**) but did not associate with expansion of CAZyme, protease or effector gene repertoires (**Supplementary Fig. 6**). Comparative genomics across the 69 *P. cucumerina* isolates using 15,269 orthogroups identified a large core genome (> 95% of genes detected in 80-100% of the strains) and a modest pan genome (< 5% of the genes detected in less than 80% of the strains) (**Supplementary Fig. 7a,b**). Pan-genome encoded genes were enriched in candidate effectors (**Supplementary Fig. 7c**) and more closely associated with transposons compared to core genome-encoded genes (**Supplementary Fig. 7d**). Between-strain differentiation in CAZyme, protease and effector repertoires was extensively explained by the *Plectosphaerella* phylogeny (54%, 36%, and 45%, respectively, PERMANOVA, *P* < 0.001, **Fig. 2c**) but not by the phylogeny of the host of isolation (*P* > 0.05) (**Fig. 2d**), thereby corroborating that the origin of the host plant is not a strong driver of repertoire variation in *P. cucumerina.* Similar results were observed for orthogroups (**Supplementary Fig. 8a,b**). We next tested whether these genetic variations associate with phenotypic differentiation for two major traits, namely 1) effect on *A. thaliana* development 28 days post inoculation in a gnotobiotic agar-based plate system (**Fig. 2b** and **Supplementary Fig. 9**) and 2) carbon resource consumption using PM1-2 Biolog® Plates that contained 190 carbon sources (**Supplementary Fig. 10a**). Although these strains were isolated from markedly different environments worldwide, they displayed little phenotypic variation for these traits. Most *P. cucumerina* isolates were detrimental on Arabidopsis, irrespective of their hosts of isolation (ANOVA *P* = 2e-16, Dunnett’s post-hoc test vs. mock, *P* < 0.05 for 44/69 *P. cucumerina* strains), contrasting with *P. plurivora* and *P. delsorboi* isolates that were non-pathogenic under these conditions (**Fig. 2b** and **Supplementary Fig. 9**). These isolates also showed remarkably similar growth phenotypes when inoculated on the 190 metabolites used as sole carbon sources (coefficient of variation < 6.5%, **Supplementary Fig. 10a,b**), and are well adapted to efficiently use plant cell wall-derived monosaccharides as growth substrates (**Supplementary Fig. 10a)**. The data suggest modest genetic variation across *P. cucumerina* isolates that is unlikely the sole factor driving adaptation to a broad range of environmental and host conditions worldwide.

**Fig. 2:**
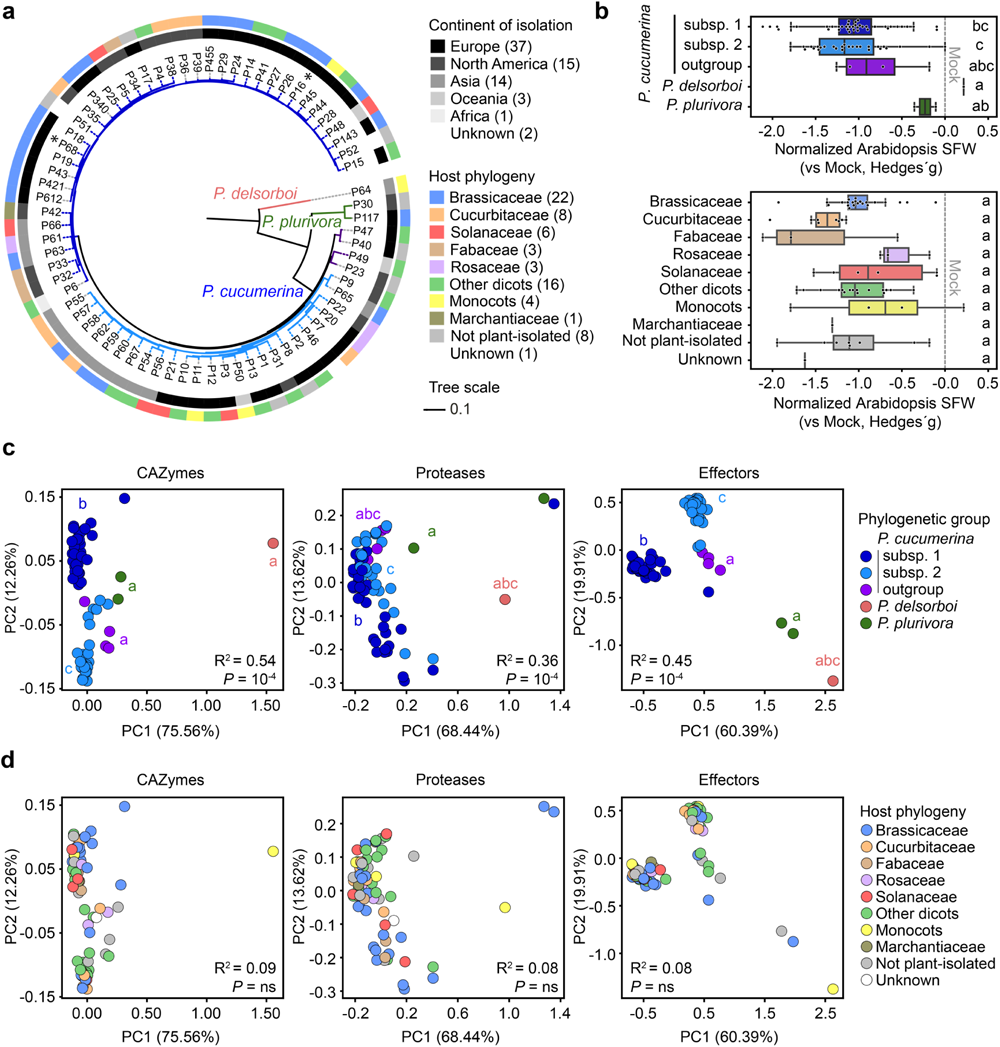
*P. cucumerina* strains isolated from phylogenetically diverse plants do not show host preference signature. **a**, Phylogeny of the 72 *Plectosphaerella* isolates, annotated with their continent of isolation (inner circle) and the phylogenetic group of their host (outer circle). The phylogenetic tree was reconstructed from the concatenated alignments of 5,466 single copy orthologs (see Methods). Two *P. cucumerina* subspecies can be distinguished and are depicted in two shades of blue whereas four strains do not belong to any of these subspecies and are depicted in violet (later referred to as “outgroup”). Isolates P68 and P16 that respectively correspond to *P. cucumerina* BMM (PcBMM, see previous work^39^) and *P. cucumerina* 0016 (see this work) are highlighted with an asterisk. **b**, Individual fungal effects on *A. thaliana* Col-0 plants organized according to the phylogenetic group of *Plectosphaerella* strains (top), or family names of the hosts of isolation (bottom). Standard effect sizes (Hedges’ g) were calculated to compare the shoot fresh weights of 28-day old inoculated and mock-treated plants (agar-based gnotobiotic system, n = 12-20 plants per fungal conditions). Letters describe differences between categories according to a ANOVA-TukeyHSD post-hoc test, performed after ANOVA testing. **c-d**, Principal Coordinate Analyses profiling inter-strain differences in genomic repertoires of CAZymes, proteases and candidate effectors across all *Plectosphaerella* isolates (n = 72 strains). The PCoA plots are color-coded according to *Plectosphaerella* phylogenetic group (**c**) or family names of the hosts of isolation (**d**). R^2^ and *P* resulting from PERMANOVA tests assessing the effect of these two categories on the three genomic repertoires are shown for each graph. Letters indicate significant differences between groups based on pairwise permutation MANOVAs followed by *P* value adjustment using the Benjamini-Hochberg method.

### *P. cucumerina* 0016 colonizes Arabidopsis, tomato and barley roots but only causes disease in dicot hosts

We next asked whether a single strain initially isolated from *A. thaliana* roots (P16, hereafter named *P. cucumerina* 0016, subspecies 1) is able to colonize/cause disease in tomato (*Solanum lycopersicum cv.* Micro-Tom) and barley (*Hordeum vulgare*) which diverged from *A. thaliana* around 110, and 140 MyA, respectively^45, 46^. Re-colonization experiments in a sterile vermiculite matrix using *P. cucumerina* 0016 and germ-free Arabidopsis, tomato, and barley seedlings (see methods) revealed contrasting effects on plant health and survival 21 days post fungal inoculation (i.e., Plant Performance Index: PPI = survival rate x shoot fresh weight, **Fig. 3a**). Whilst the fungus was highly detrimental on both tested dicotyledonous plants (19.5% and 10.7% reduction in PPI for Arabidopsis and tomato, respectively) (two-sided t-test or Wilcoxon rank-sum test, *adj. P* < 0.05), it remained harmless when inoculated on barley roots (*adj. P* > 0.05, **Fig. 3a**). Inspection of fungal load in roots of these plants using quantitative PCR (qPCR) indicated that the fungus was able to colonize all roots, including those of barley (two-sided t-test or Wilcoxon rank-sum test, *adj. P* < 0.05, **Fig. 3b**). We next stained infected roots using calcofluor white and WGA CF®633A for visualizing the spatial distribution of fungal hyphae during infection. Consistent with our amplicon sequencing data (**Fig. 1a**), numerous *P. cucumerina* 0016 extraradical hyphae were observed throughout the entire root system of Arabidopsis, from the mature peridermal region to the root tip, including lateral roots (**Fig. 3c**). Penetration of undifferentiated hyphae was repetitively observed into or between peridermal (**Fig. 3c-i, iii, iv**) or cortical cell layers (**Fig. 3c-ii, v**), thereby validating extensive endophytic propagation (**Fig. 3c**). In tomato roots, epidermal walls were coated with extracellular hyphae and endophytic colonization was primarily confined to the first epidermal cell layer (**Fig. 3d**). In contrast, despite extensive fungal proliferation at the vicinity of barley root hairs, we did not observe endophytic proliferation in barley roots, suggesting that the fungus is unable to invade and/or is stopped by the barley immune system (**Fig. 3e**). Taken together, our data suggest host-specific differences in fungal endophytic colonization that correlate with distinct outcomes on host performance (**Fig. 3a**).

**Fig. 3:**
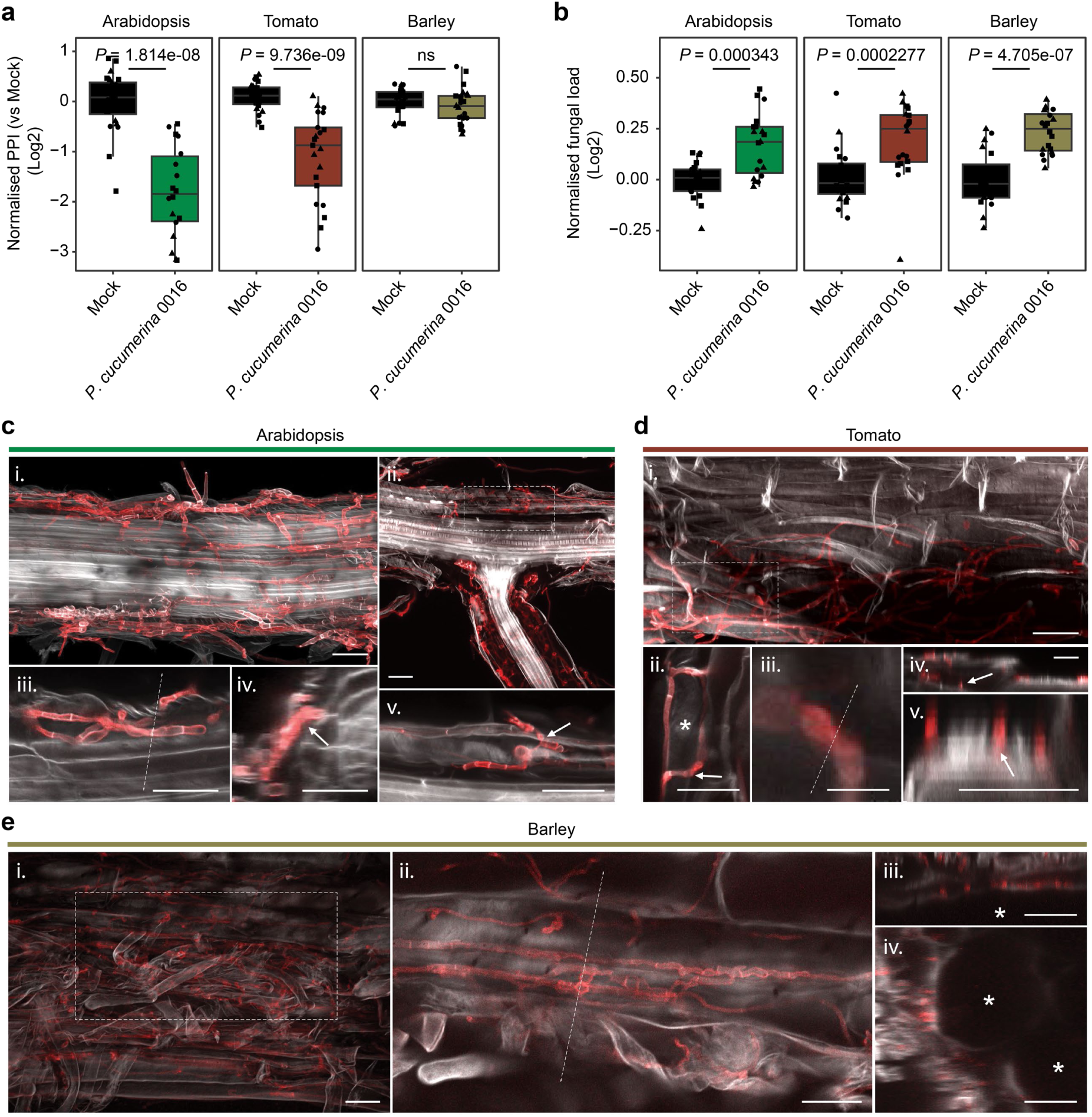
Colonization of *P. cucumerina* 0016 in Arabidopsis, tomato and barley roots and impact on host performance: **a**, Plant Performance Indices (PPI) of Arabidopsis, tomato, and barley colonized by *P. cucumerina* 0016 were normalized to mock-treated plants and log2 transformed. PPI values were calculated by multiplying the survival rate by the shoot fresh weight of the plants. **b**, Relative abundance of *P. cucumerina* 0016 in the roots of Arabidopsis, tomato, and barley measured by qPCR-based quantification of the fungal ITS *vs* plant *UBQ* gene. **a**-**b**, Pre-grown seedlings (2-week-old Arabidopsis and tomato and a week-old barley) were inoculated with *P. cucumerina* 0016 and harvested 21 days post fungal inoculation (vermiculite-based gnotobiotic system). Three independent biological replicates are depicted (circles, triangles, squares), each consisting of 4 – 8 plants (n = 12 – 24 plants per condition). Pairwise comparisons of plant performance between mock-treated and *P. cucumerina* 0016 treated plants were performed using either the two-sided t-test or Wilcoxon rank-sum test (*P* ≤ 0.05). **c**-**e**, Confocal imaging of Arabidopsis (**c**), tomato (**d**), barley (**e**) roots colonized by *P. cucumerina* 0016 21 days post fungal inoculation. Roots were double-stained with Calcoflour white and Wheat Germ Agglutinin coupled to fluorophore CF633A (WGACF633; Biotium) and imaged using Zeiss LSM 880 confocal microscopy. **c**, i. Maximum intensity projection of Z-stack images of an Arabidopsis primary root extensively colonized. ii. Maximum intensity projection of middle section of the primary root from which a lateral root emerges. iii. Enlargement of one optical section from (i), iv. Orthogonal projection of (iii) showing undifferentiated hyphae penetrating peridermal cell layer (arrow). v. Enlargement of one optical section shown in (ii) (dashed square) depicting an intracellular hypha colonizing the root cortex (arrow). **d**, i. Maximum intensity projection of a middle section of a primary root. ii. Enlargement and Z-stack image of (i) (dashed square) showing intercellular hyphae (arrow) surrounding an epidermal cell (star). iii. Enlargement of one optical section shown in (ii) (arrow). iv and v. Orthogonal projection of (ii). Undifferentiated hyphae adhere to the epidermal cell (arrows). **e**, i. Maximum intensity projection of the middle section of the root. ii. Z-stack slice of (i) (dashed square) showing the colonization of fungal hyphae in the epidermal layer of the barley root. iii and iv. Orthogonal projection of (ii) showing hyphae adhering to epidermal cells (stars).

### *P. cucumerina* 0016 transcriptional reprogramming *in planta* is modulated by host identity

Given the low intra-species genomic variation observed in *P. cucumerina*, we hypothesized that fungal adaptation to a broad range of hosts is conferred by activation of common gene sets driving “general” root infection, but also specific subsets responding to variation in host physical and chemical cues. Using the same vermiculite system described above, we sequenced the *P. cucumerina* 0016 transcriptome 21 days post inoculation on Arabidopsis, tomato, and barley roots and compared this *in planta* transcriptome to that of a control fungal transcriptome obtained from inoculated polyester membranes (Ctrl, *ex planta*, see Methods). Mapping of Illumina short reads against corresponding reference genomes (**Supplementary Fig. 11a,b**), followed by statistical analysis of differentially expressed genes (DEGs*, in planta* vs Ctrl, DESeq2, Generalized Linear Model) identified 3,399 fungal genes significantly regulated in response to the different hosts (**Supplementary Table 4**). Multidimensional scaling (MDS) of Pearson distances computed on normalized read counts of both expressed genes (**Supplementary Fig. 12**) and DEGs (**Fig. 4a**) identified fungal transcriptome differentiation that is primarily shaped by *in planta* vs *ex planta* conditions (i.e., see x-axis explaining 60.85% of the variation, **Fig. 4a**), but also to a major extent by the host plant identity (i.e., see y-axis explaining 30.63% of the variation; pairwise statistical analyses: ANOVA-TukeyHSD, *adj. P* < 0.05, **Fig. 4a**). Comparison of all fungal DEGs identified for each of the three comparisons (n = 1,862 induced, n = 1,537 repressed in total, *in planta* vs Ctrl) revealed that 19.3% and 37.6% were consistently induced and repressed, respectively, irrespective of the host plant (ATB sector, **Fig. 4b,c**). Consequently, most fungal DEGs responded either to two plant species (AT, AB and TB sectors, 22.0% induced, 20.7% repressed; **Fig. 4b,c**) or were host-specific (A, T and B sectors: induced: 58.7%; repressed: 41.7%; **Fig. 4b,c**), thereby illustrating high transcriptome plasticity in response to different hosts (**Fig. 4d**). Notably host-specific DEGs prevailed during infection on Arabidopsis (A sector) compared to tomato (T sector) or barley (B sector), potentially reflecting higher adaptation to the host of isolation (**Fig. 4c,d**). GO-term enrichment analysis (*adj. P* < 0.05; Benjamini-Hochberg method) for each of the above-mentioned DEG sectors (**Fig. 4c**) revealed remarkable enrichment of terms related to catabolism that were exclusively observed in the up-regulated gene sets (**Fig. 4e** and **Supplementary Fig. 13a,b**). Those included: “Hydrolase”, “Proteolysis”, “Cellulase”, “Catalytic activity”, “Pectate lyase” or “Subtilase”, that were often identified as enriched in both host-specific (A, T, B) and shared (ATB) fungal DEGs subsets. Taken together, our results revealed that *P. cucumerina* transcriptional reprogramming is differently tailored depending on the colonized host and that catabolism of host-derived products is likely key in driving infection across a broad range of hosts.

**Fig. 4:**
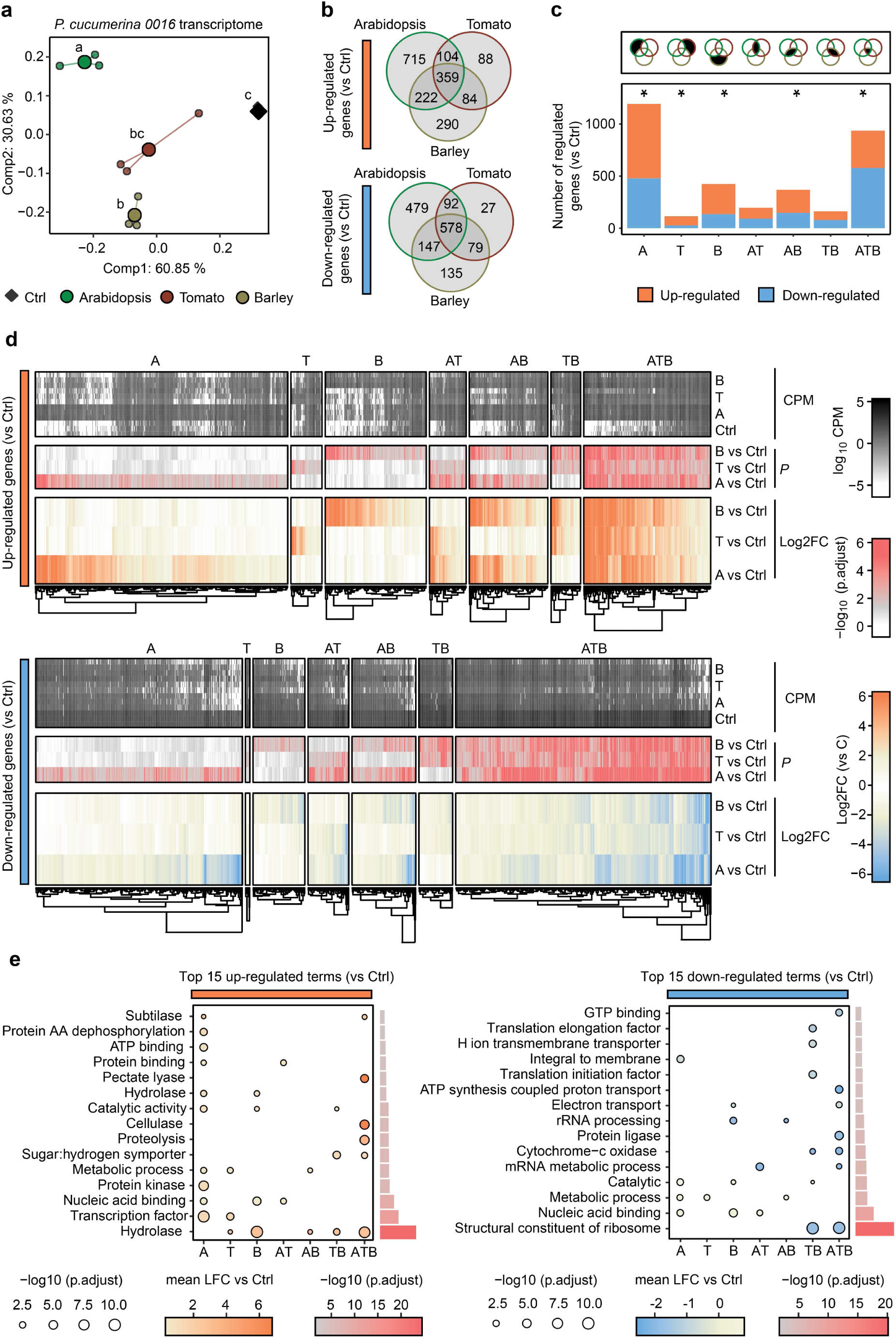
*In planta* transcriptome of P. *cucumerina* 0016 is modulated by host plant identity. **a**, Multidimensional Scaling (MDS) plot of Pearson distances computed on normalized read counts of differentially expressed genes (DEGs, n = 3,399, *in planta* vs *ex planta*). The different plant hosts are shown in green (Arabidopsis), red (tomato), and khaki (barley). *Ex planta* control condition is depicted in black (Ctrl, membrane). Large data points represent the mean values of the respective datasets in MDS1 and MDS2, whereas small data points represent the three biological replicates. Multiple statistical comparisons across conditions were performed using TukeyHSD test (*P* ≤ 0.05). **b**, Venn diagram illustrating the number of DEGs (|log2FC| ≥ 1 and FDR ≤ 0.05) that are up-regulated (orange) and down-regulated (blue) *in planta vs* Ctrl condition. **c**, Stacked bar graph of fungal DEGs that are up- and down-regulated *in planta.* The categories A, T, and B denote fungal genes uniquely up- or down-regulated in Arabidopsis, tomato, or barley roots, respectively. AT, AB, and TB denote genes commonly up- or down-regulated in two hosts (Arabidopsis and tomato, Arabidopsis and barley, or tomato and barley, respectively), while ATB denotes genes consistently up- or down-regulated across all hosts. Categories that are enriched in up- or down-regulated DEGs *in-planta* are marked with an asterisk (*, proportion test with Pearson’s chi-square). **d**, Heatmaps depicting DEGs that are up-regulated (orange) and down-regulated (blue), respectively. Bottom heatmaps depict log2FC of DEGs that are hierarchically clustered along the y-axis, with corresponding conditions displayed along the x-axis. Middle heatmaps depict *adj. P* of DEGs (Benjamini-Hochberg correction, -log10). Top heatmaps show counts per million of DEGs (log10 scale). **e**, Gene Ontology (GO) term enrichment analysis. Top 15 significantly enriched GO terms that are up (left) and down regulated (right), respectively, *in planta vs* Ctrl. The x-axis represents A, T, B, AT, AB, TB, and ATB sectors (see above) and the y-axis the GO terms. The color of each sphere represents the mean log2FC of DEGs, while the size of the sphere indicates the mean *adj. P* within each GO category for the respective conditions. The bar graph illustrates the cumulative *adj. P* for the respective GO. These values were computed using the hypergeometric test with Benjamini-Hochberg correction (*P* ≤ 0.05).

### Activation of CAZyme-encoding genes prevails during infection and mirrors root cell wall composition

We next hypothesized that *P. cucumerina* 0016-mediated cell wall degradation prevails during root colonization and plastically adjust according to host cell wall polysaccharide composition. Consistent with our GO-term enrichment analysis (**Fig. 4e**), we observed that genes encoding carbohydrate active enzymes (CAZymes) were disproportionally upregulated during root infection in all above-mentioned DEG sectors (A: 78/715 = 10.9%, T: 19/88 = 21.6%, B: 59/290 = 20.3%, AT: 17/104 = 16.3%, AB: 44/222 = 19.8%, TB: 17/84 = 20.2%, ATB: 101/359 = 28.1% of the DEGs, **Supplementary Fig. 14a-c**), thereby contrasting with the expected proportion of CAZyme genes encoded in *P. cucumerina* 0016 genome (9.9%). Next, we constructed a force-directed network map (see methods), where circular nodes represented plant cell wall-degrading CAZyme families (n = 83, see methods) connected to their predicted plant substrates (as registered in the dbCAN-sub database^47, 48^ (n = 14, see methods). By mapping the mean Log2FC of CAZyme families significantly overexpressed *in planta* (see circle size and grey gradient, Log2FC, Fisher’s-exact test, *P* < 0.05) onto the network backbone for each above-mentioned DEG sectors, we predicted host cell wall components (square nodes, orange gradient, Log2FC, Fisher’s-exact test, *P* < 0.05) that were targeted by the fungus in response to a specific host (A, T, B, **Fig. 5a**), two hosts (AT, AB, TB, **Fig. 5a**) or all three hosts (ATB **Fig. 5a,b**). This revealed that diverse CAZymes were predicted to target all major cell wall constituents (pectic polysaccharides, cellulose, hemicellulose, lignin), irrespective of the host plant (ATB sector, **Fig. 5a-c**). These CAZymes belong to polysaccharide lyase (PL) families 1_7, 3_2, 9_3, and 42, the carbohydrate binding module family 32, the glycoside hydrolase (GH) family 64, lytic polysaccharide monooxygenases families AA9, AA16, and the carbohydrate esterase family 2 (**Fig. 5b**). Cell walls of dicotyledonous plants are more pectin-rich than those of monocots, while monocot walls contain mainly structurally divers xylans and mixed-linkage glucans as their dominant hemicelluloses^49^.This is consistent with our targeted analysis of wall materials in roots of Arabidopsis, tomato, and barley as pectic uronic acids were more prevalent in Arabidopsis and tomato, while (arabino)-xylan and matrix polysaccharide based glucose were more prevalent in barley (**Supplementary Fig. 15**). Interestingly, these differences correlated with host-specific activation of fungal genes encoding pectin-degrading enzymes in the dicot hosts (DEG sectors A, T, AT; PL3_2, PL 1_7, PL11_2 families), or (arabino)xylan-degrading enzymes in the barley host (DEG sector B; GH51_2) (**Fig. 5a,c**). Our results suggest that sensing and adequate transcriptional response to cell wall physicochemical cues contributes to plastic infection of diverse hosts in *P. cucumerina*.

**Fig. 5:**
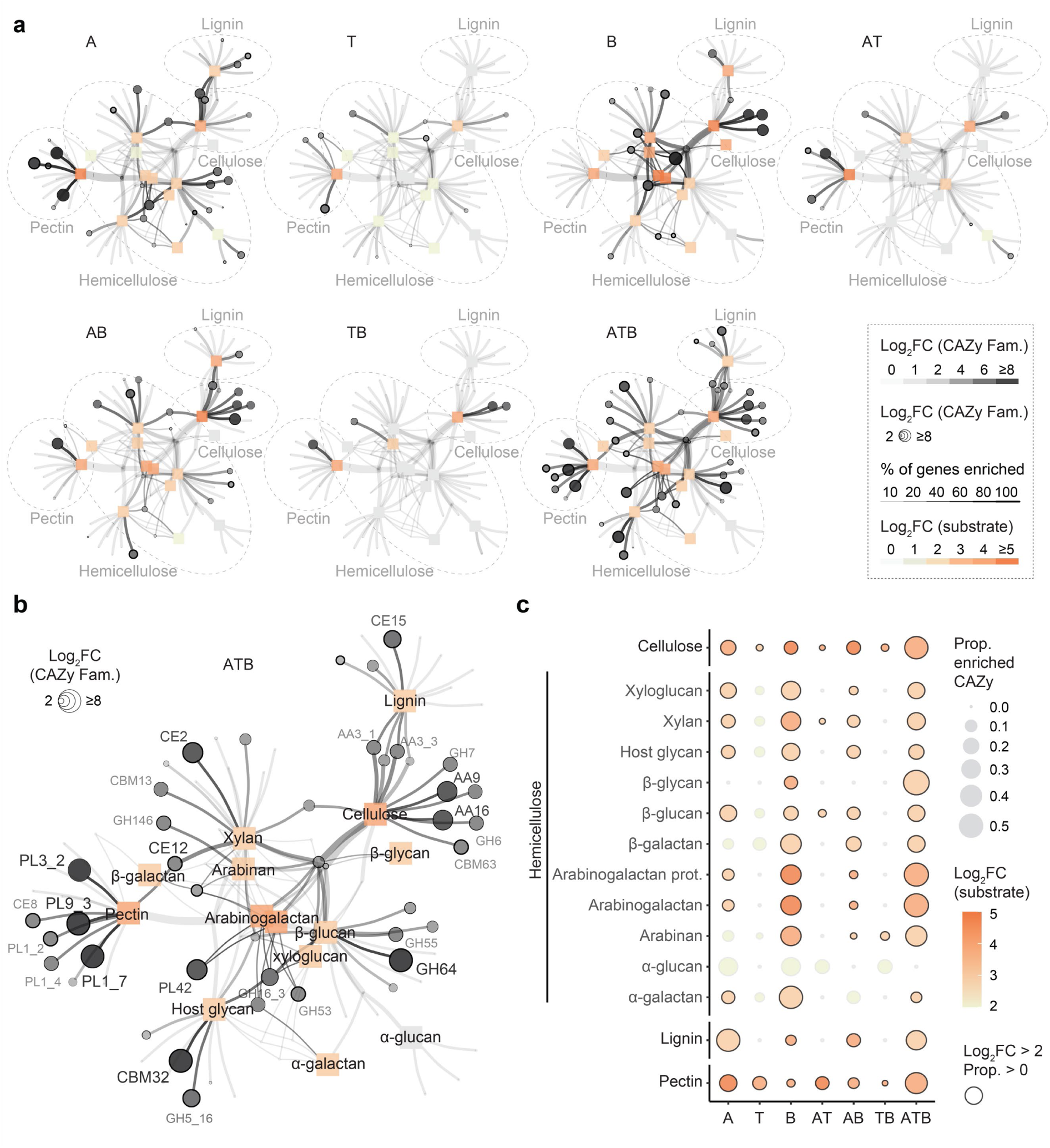
Host-specific and core sets of *P. cucumerina* 0016 CAZyme-encoding genes induced during root colonisation of different hosts. **a-b,** The networks illustrate CAZyme families acting on plant and plant/fungal cell walls (circles) alongside with their potential plant cell wall substrates (n = 14 substrates, squares) defined based on the CAZyDB^119^ and dbCAN-sub databases^47^. The color gradient of the nodes and edges (white to black) and the size of the circles reflect the mean Log2FC of statistically enriched CAZyme family *in planta* vs *ex planta* (Fisher’s exact test) for each DEG subset (**a,** A, T, B, AT, AB, TB, ATB **b**, enlargement of the ATB network, see **Fig. 4c**). For each CAZyme family, the thickness of the circles indicates the proportion of genes induced among the DEGs across conditions. Predicted substrates targeted by CAZymes are shown using a grey-to-orange gradient that depicts the mean Log2FC of the enriched CAZymes targeting specific substrates across conditions. Substrates represented in grey squares are classified as non-significant (with a proportion of enriched CAZymes equals to 0). **c,** Balloon plot illustrating the utilization of substrates through the differential expression of fungal CAZymes based on their log2FC *in planta* vs *ex planta*. The colors within the circles (yellow-to-orange) depicts the mean log2FC of the CAZymes predicted to target primary cell wall substrates such as cellulose, diverse hemicellulose components, lignin, or pectin (see y-axis) for the different DEGs subsets (see x-axis, A, T, B, AT, AB, TB, and ATB). The size of the circles corresponds to the proportion of DEGs within the CAZyme family that target the different cell wall substrates (y-axis). Circles marked in black represent groups that show a mean log2FC along with a proportion of enriched CAZymes > 0 and in grey where the proportion of enriched CAZymes equal to 0, as determined by Fisher’s exact test.

### GH64 CAZyme-encoding gene promotes multi-host infection and disease in *P. cucumerina* 0016

Based on our network map (ATB sector, **Fig. 5b**), we identified a single copy gene encoding a secreted GH64 enzyme (jgi|Plecuc1|282412|) that displayed one of the highest log2FC enrichment *in planta*, irrespective of the host plants (Arabidopsis: Log2FC = 7.4, tomato: Log2FC = 8.3 barley: Log2FC = 8.3, **Supplementary Table 4**). Following AlphaFold protein structure prediction, a FoldSeek^50^ search identified strong structural similarity with the SpGlu64A enzyme secreted by *Streptomyces pratensis A* (**Fig. 6a, Supplementary Fig. 16a**), a β-glucanase exhibiting strict substrate specificity toward β-1,3-glucans^51^. These β-1,3-linked glucose units are constituents of plant cell walls (i.e. callose and grass-specific hemicellulosic mixed-linkage glucans), but mainly represent immunogenic molecules of fungal cell walls^52, 53, 54, 55^. By transfecting *P. cucumerina* 0016 protoplasts with a ribonucleoprotein (RNP) complex consisting of the Cas9 protein and a single guide RNA (sgRNA), we drove precise insertion of a phleomycin resistance DNA cassette into the coding sequence of *GH64* via homologous recombination (method, **Supplementary Fig. 17a-d**). Corresponding mutant spores inoculated on germ-free roots of Arabidopsis and tomato seedlings showed 62%-77% reduction in pathogenicity (Kruskal-Wallis-Dunn Test, adjusted with Bonferroni correction*, P* < 0.0001, **Fig. 6b**) and 18%-24% reduction in root colonization in tomato and Arabidopsis plants (Kruskal-Wallis-Dunn Test, adjusted with Bonferroni correction*, P* < 0.0001, **Fig. 6c**) compared to the WT strain. In contrast, no significant differences were observed in the barley host (*P* > 0.05, **Fig. 6b,c**), nor in artificial growth media (**Fig. 6d**, **Supplementary Fig. 17e,f**), indicating that the gene is a root colonization determinant of the two tested dicot hosts. WT-like disease phenotypes were also observed for another mutant (*Δpl11_2*, **Supplementary Fig. 18**), excluding the possibility that phleomycin resistance is influencing these phenotypes. Notably, GH64 shows high sequence conservation across the 72 *Plectosphaerella* isolates (**Supplementary Fig. 16b**), has orthologs in specific fungal lineages belonging to Sordariomycetes and Dothideomycetes and is significantly enriched in genomes of plant saprotrophs, pathogens and endophytes (n = 2,534 fungal genomes, **Fig. 6e** and **Supplementary Fig. 16c,d**). Remarkably, inactivation of the corresponding GH64 orthologous gene in the root pathogen *Colletotrichum incanum*^4, 34, 56^ (class Sordariomycetes, family Glomerellaceae, jgi|Colin1|11312|, **Supplementary Fig. 19a**) resulted in 68.5% reduction of fungal mutant load in *A. thaliana* roots and 10% increase in shoot fresh weight compared to the WT *C. incanum* strain (24 days post infection, **Fig. 6f,g**). However, no significant differences were noted in the Solanaceae host *Nicotiana benthmiana* (i.e., tobacco **Supplementary Fig. 19b-d**). Taken together, our data indicate that *in planta* activation of this gene in evolutionary-distant fungal lineages promotes root infection and disease.

**Fig. 6.**
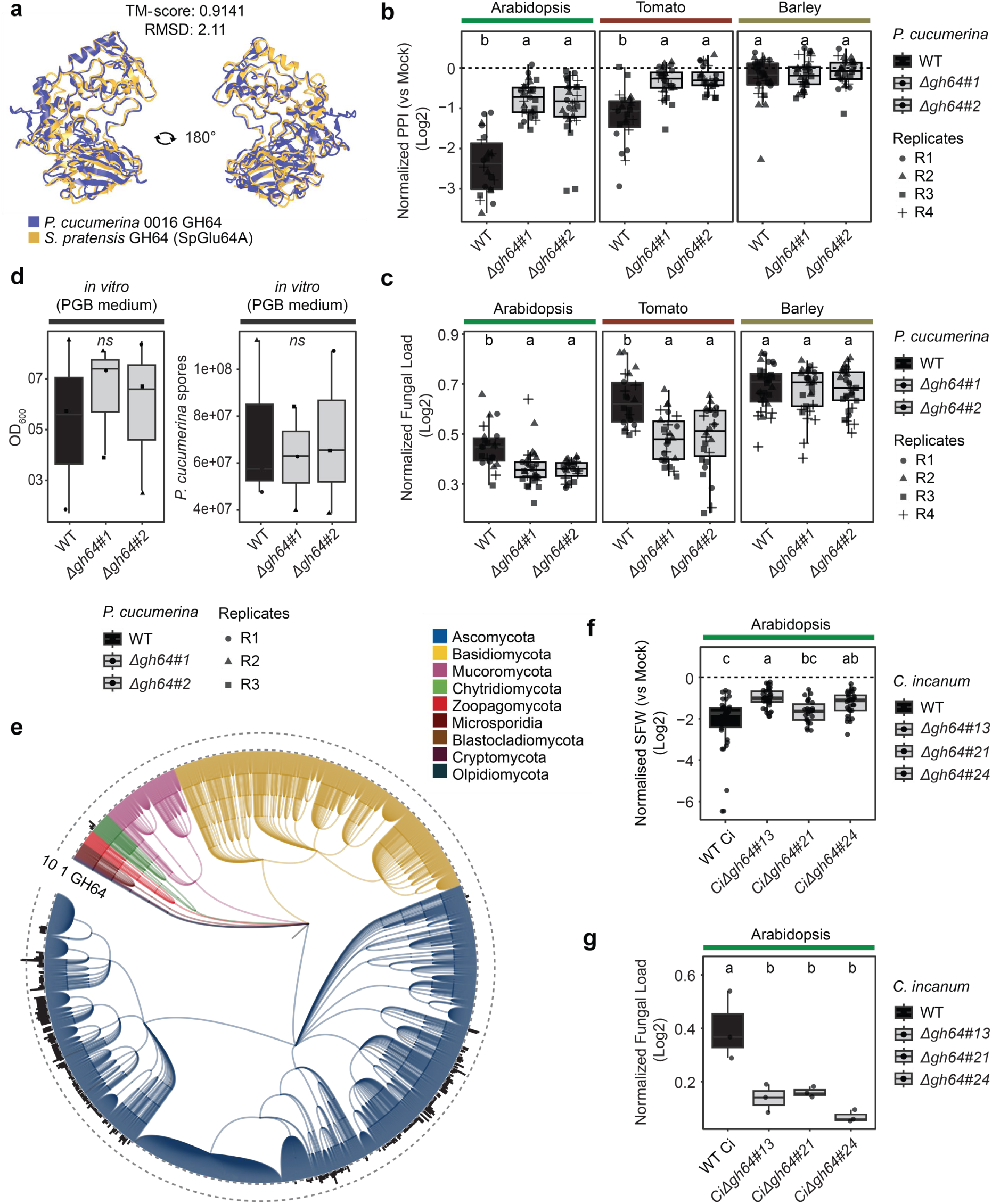
Phenotype assessment of *P. cucumerina* 0016 *and C. incanum Δgh64* mutant strains: **a,** AlphaFold3-predicted structure^132^ of *P. cucumerina* 0016 GH64 (in blue; pTM=0.92) showing high similarity to the previously resolved structure of the GH64 enzyme SpGlu64A of *Streptomyces pratensis*^51^ (in orange), according to FoldSeek search analysis^50^. **b,** Plant Performance Indices (PPI, see method) of Arabidopsis, tomato, and barley colonized by *P. cucumerina* 0016 were normalized to mock-treated plants and log2 transformed. PPI values were calculated by multiplying the germination rate by the shoot fresh weight of the plants. **c**, Relative abundance of *P. cucumerina* 0016 in the roots of Arabidopsis, tomato, and barley measured by qPCR-based detection of the fungal ITS vs. plant *UBQ10* gene. **b-c**, Pre-grown seedlings (2-week-old Arabidopsis and tomato and a week-old barley) were inoculated with *P. cucumerina* 0016 and harvested 21 days post fungal inoculation. Four independent biological replicates are depicted (circles, triangles, squares, and plus), each consisting of 5 – 8 plants (n = 20 – 32 plants per condition). Multiple pairwise comparisons of plant performance between mock-treated and *P. cucumerina* 0016 treated plants were performed using Kruskal-Wallis-Dunn Test, adjusted with Bonferroni correction (*P* ≤ 0.05). **d,** In vitro growth assessment of *P. cucumerina* 0016 WT and *Δgh64* mutant strains. An initial spore inoculum of 8x10^4 spores/mL was used and the fungus was inoculated on PGB medium for 24 hours. OD_600_ was measured after 24 hours using a spectrophotometer (left) and yield of *P. cucumerina* 0016 spores were counted after 24 hours using a hemocytometer. **e,** Phylogenetic tree constructed using 2,534 fungal genomes from the MycoCosm^85^ (JGI) with NCBI taxonomy indices. The colors of the clade indicate the fungal phylum. The barplot at the leaf nodes denotes the number of GH64 occurrences that are identified in the genomes (CAZyme family-GH64 from JGI mycocosm). **f,** Shoot fresh weight (SFW) of *A. thaliana* plants colonized by *Colletotrichum incanum* wild type (WT *Ci*) and three independent mutant lines (*CiΔgh64#13*, *#21*, *#24*) normalized to mock-treated plants (dashed line) and log2 transformed. Plants (n = 26 - 36 per condition) were harvested at 24 dpi. Statistical significance was determined by ANOVA-TukeyHSD, *P* < 0.05. **g,** Relative load of WT *Ci* and mutant strains in the roots of *A. thaliana* measured by qRT-PCR-based detection of *C. incanum* ACTIN transcripts normalized to Arabidopsis ACTIN transcripts at 24 dpi (n = 3 replicates per conditions, 8 - 12 plants were pooled per replicate). Statistical significance was determined by ANOVA-TukeyHSD, *P* < 0.05.

## Discussion

Our study reveals the remarkable occurrence of *P. cucumerina* in the roots of diverse host plants, including the model plant species *A. thaliana*. *P. cucumerina* therefore represents one of the very few core members of the Arabidopsis root mycobiota, based on both culture-dependent and independent approaches. Despite this, our analysis revealed that *P. cucumerina* isolates are reproducibly causing disease when inoculated on germ-free *A. thaliana* roots, irrespective of their hosts of isolation. We previously showed that the most prevalent fungal isolates colonizing *A. thaliana* roots in nature are often harmful in mono-association experiments^8^ and that these detrimental phenotypes are alleviated by the presence of bacterial commensals that regulate fungal load and composition at the root interface, thereby preventing fungal dysbiosis in nature^9, 10, 12^. It is therefore conceivable that *P. cucumerina*-induced disease in nature relies on both the genetic determinants identified here for multi-host infection but also on other genetic factors driving root microbiome manipulation, as previously reported for closely related *Verticillium* soil-borne pathogens^57, 58, 59^.

Our results indicate that the ability of *P. cucumerina* to colonise multiple hosts across various environments is at least to some extent mediated by a remarkably large repertoire of CAZymes acting on diverse plant cell wall substrates. In comparison with 93 other plant-associated Sordariomycete fungi^8^, *P. cucumerina* 0016 had a unusually diverse repertoire of plant cell wall degrading enzymes (PCWDEs, 12^th^/94, n = 160, range 198-29), especially those acting on cellulose (3^rd^/94, n = 92, range 94-6) and displayed the highest percentage of PCWDEs within fungal CAZyme repertoires (1^st^/94, 76%, range 76%-33%)^8^. Our within-species comparative genome analysis amongst 69 *P. cucumerina* isolates revealed surprisingly little variation in gene repertoire categories, including CAZymes (range 1,212-1,397, median = 1,291), which did not correlate with variation in fungal detrimental activity. We propose that among-strain phenotypic variation in pathogenicity arose from differences in gene expression, rather than in gene repertoire composition, as previously shown for closely related fungal isolates with contrasted lifestyles^30, 34, 60, 61^.

We also report the capacity of *P. cucumerina* to express differential sets of CAZymes upon sensing of host-derived cues as a potential strategy driving multi-host compatibility. Indeed, root-infecting fungi colonizing both dicot and monocot hosts have likely evolved mechanisms to efficiently respond to markedly different root cell walls. Such sensing and adequate transcriptional activation of fungal genes encoding PCWDEs was previously reported for broad host range leaf pathogens^62, 63^, and our data support that fungal sensing of host-specific cell wall cues might contribute to host range expansion belowground. Therefore, *P. cucumerina* dispersion success in nature is likely explained by its capability to infect multiple hosts at any sites in order to avoid population bottleneck. Our results are consistent with the idea that widespread soil-borne fungal pathogens often have a broad host range and remarkably diverse CAZyme repertoires. Therefore, we conclude that both diverse and tuneable repertoires of PCWDEs promotes multi-host infection.

We showed that infection of *A. thaliana* and *S. lycopersicum* roots by *P. cucumerina* requires a candidate β-1,3 glucanase GH64, thereby promoting disease susceptibility. Given that GH64-encoding gene is induced in response to a broad range of hosts, it is plausible that the root signal is evolutionary conserved across monocots and dicots. However, because endophytic fungal colonization was limited in barley roots and that pathogenicity was observed in multiple dicot hosts that diverge at least 110 MyA (*A. thaliana* and *S. lycopersicum*), we propose that this candidate β-1,3 glucanase promotes *P. cucumerina* proliferation in the root endosphere. This is consistent with previous observations showing that endophytic colonization aggressiveness and disease symptoms are often linked during root colonization by mycobiota members^7, 8^. The differential enrichment of this gene in genomes of saprotrophs, endophytes and pathogens also suggests a broader relevance for plant cell wall penetration and/or degradation in diverse fungal lineages. Remarkably, mutation of the orthologous gene in the root pathogen *C. incanum* also prevented root infection and disease in *A. thaliana*. However, the gene was dispensable for tobacco root infection, consistent with its low expression in the roots of this host. Irrespective of this, our data corroborate that GH64 is a root infection determinant in at least two fungal pathogens that evolved independently in distinct families within Sordariomycetes.

β-Glucans represent a diverse group of polysaccharides that are found in both plant and fungal cell walls^64^. These glucans can be recognized as MAMPs (Microbe-Associated Molecular Patterns) or DAMPs (Damage-Associated Molecular Patterns) and initiate plant immune responses^54, 65, 66^. To suppress β*-*glucan-mediated immune activation, plant-colonizing fungi have been shown to use various strategies^67^, including down-regulation of β-glucan biosynthesis genes *in planta*^68^, secretion of lectins that bind β-glucans^69^ or of molecules that inhibit plant endoglucanase^70^. *Magnaporthe oryzae* was recently shown to secrete a GH17 exo-β-1,3-glucanase (Ebg1) that hydrolyzes β-1,3-glucans into glucose, thereby preventing plant immune responses^21^. Similarly, a fungal insect pathogen was shown to deploy a host-inducible β-1,3-glucanase called BbEng1 in order to deplete immunogenic β-1,3-glucans from fungal cell walls, thereby evading host immune surveillance and promoting successful fungal mycosis^71^. Therefore, suppression of β-1,3-glucan-induced immune activation could represent one strategy deployed by *P. cucumerina* to promote endophytic infection in roots of Arabiodopsis and tomato. Furthermore, *P. cucumerina* could also deploy this GH64 enzyme *in planta* to break down callose, a plant cell wall polysaccharide composed of β-1,3-linked glucose units^72, 73^. Given that callose deposition is well known to occur at pathogen infection sites and in plasmodesmata to block pathogen invasion^74, 75, 76^, it is tempting to speculate that *P. cucumerina* could also have evolved this strategy to overcome callose-mediated restriction of fungal propagation in roots. Irrespective of the mode of action, GH64 represents a target of choice to efficiently control destructive root pathogens that have a broad host range.

## Methods

### Re-analysis of *A. thaliana* mycobiota profiling data

Data from a European transect study profiling the root mycobiota of *A. thaliana* at 17 sites were re-analysed^14^, together with profiling data of *A. thaliana* roots grown in Cologne agricultural soil^9^. Paired sequencing reads were joined and demultiplexed using QIIME^77^ and its scripts *join paired reads* and *split libraries fastq*. Quality filtering step was performed with phred score of 30 or higher. For fungal reads that could not be merged, the respective forward reads were kept for further processing. Filtered and demultiplexed reads were trimmed to an equal length of 220bp using USEARCH with option-fastx truncate^78^. Those reads were dereplicated and forwarded to the UNOISE3 pipeline^40^. Amplicon sequence variants (ASVs) were determined using the USEARCH unoise3 command, where reads with sequencing errors are corrected and possible chimeras are removed, leaving a set of correct amplicon sequences. ASVs reads were checked against an ITS sequences database (full-length ITS sequences from the National Center for Biotechnology Information (NCBI)) to remove non-fungal reads. Fungal ASVs were classified using the Warcup database^79^. To receive count-tables, ASV sequences were then mapped back against the whole read dataset at a 97% similarity cut-off (using USEARCH-usearch global).

### Analysis of *P. cucumerina* niches, hosts and prevalence in soils worldwide

To identify which plant hosts *P. cucumerina* colonizes in nature, we accessed the GlobalFungi online database^43^ (in November 2024). Samples in which *P. cucumerina* (SH1002406.10FU) was detected were compared to total samples registered in the database, focusing on sample niches. Overrepresentation of a niche in samples in which *P. cucumerina* was detected was tested with a one-sided Fisher’s exact test, using the formula stats.fisher_exact(alternative=’greater’) in Scipy^80^, followed by Benjamini-Hochberg *P* value correction. To identify plant species that host *P. cucumerina* in their roots and rhizosphere, “root”, “rhizosphere” and “root+rhizosphere” samples in which *P. cucumerina* was detected were browsed and the “dominant plant species” metadata associated with each sample were collected. These species were grouped by phylogenetic classes, according to the NCBI Taxonomy database^81^. The plant phylogeny presented on Figure 1 was obtained using Taxallnomy^82^ based on NCBI Taxonomy data.

To analyse the prevalence of *P. cucumerina* in soils, previously curated ITS1-5.8S-ITS2 rDNA sequences of species^35^ were downloaded and ITS1 sequences were extracted from them using ITSx^83^. We then linked these ITS1 sequences to ASVs from the Global Mycobiome data set^84^ using blastn with a percentage of identity threshold of 100%. These ASVs were detected in 780/3,194 soil samples (relative abundance > 0). Metadata of sites at which ASVs were detected were then analysed, focusing on latitude, longitude and biome types. We tested the enrichment of specific biomes in soils containing *P. cucumerina* using a random sampling approach. We randomly sampled 780 soil samples 99,999 times, and each time counted the number of soil samples associated to each biome. We used a Fisher’s exact test to compare whether the set of soils in which *P. cucumerina* was detected contains significantly more/less samples associated to each biome than the randomly sampled sets of soils. Statistical testing was performed with Scipy 1.5.3 and the function stats.fisher_exact(alternative=’two-sided’)^80^.

### Assembly of a culture collection of *Plectosphaerella* isolates

We previously isolated 7 strains of *P. cucumerina* and 1 strain of *P. plurivora* from surface-sterilized *A. thaliana* roots sampled across Europe^9^. We imported 64 additional strains from 17 collaborators or culture collections, sampled from different hosts and environments across different continents. See **Supplementary Table 3** for details.

### *A. thaliana* recolonization experiments with spores of *Plectosphaerella* isolates

We performed plant recolonization experiments with individual *Plectosphaerella* strains in an agar-based system. First, to prepare fungal spore stock solutions, fungi were grown on oatmeal medium. This medium was prepared by crushing 5 g of oatmeal in liquid nitrogen with a mortar and pestle until obtaining a powder. This powder was boiled for 30 min in 200 mL of water on a heating plate at 180°C. Then, 4 g of Difco Agar (ref. 214530) was added to the mixture prior to autoclaving. Fungi were revived from 30% glycerol stocks and grown on this medium for 10-14 days. Sterile water was poured on the grown mycelium, pipetted then filtered on Miracloth, then centrifuged at 4000 x g for 10min to concentrate spores. Spores were counted on a Malassez cell, and their concentration was adjusted to 10^7^ spores/mL in 25% glycerol. These stock solutions were flash-frozen in liquid nitrogen, then kept at -80°C. *A. thaliana* Col-0 seeds were sterilized 15 min in 70% ethanol, then 5 min in 8% sodium hypochlorite. They were then washed six times in sterile double-distilled water and one time in 10 mM MgCl_2_. They were kept 5 days at 4°C in the dark for stratification. Culture medium was prepared by mixing 2.2 g Murashige and Skooge (Duchefa Biochemie, ref. M0222) and 0.5 g MES buffer (Roth, ref. 4256.5) powders in 1 L of double-distilled water. Then, pH was adjusted to 5.7, and 10 g of Difco Agar (ref. 214530, 1% final concentration) was added. After autoclaving, 120x120mm petri dishes were filled each with 65 mL of medium. The top 2 cm of agar were cut out, and 7 seeds were deposited where the cut was made. Plates were sealed with micropore tape and their bottoms were wrapped (up to the seeds level) in paper to hide roots from the light. Plants were grown for 10 days at 21°C (10h with light intensity 4 at 19°C, and 14 h in the dark) in growth chambers. After 10 days, culture plates were re-opened and only the four biggest plants in each plate were kept. To prepare a spore inoculum, 5 μL of defrosted spore stock solution was pipetted and mixed with 995 μL of sterile water. This mixture was centrifuged at 9,000 g for 10 min. The top 950μL in the tube were replaced with fresh sterile water. With this procedure, glycerol was removed and the concentration of spores in the tube was set to 500 spores/10 μL. Ten-days old seedlings were inoculated by pipetting down 10 μL of inoculum (500 spores) on their primary roots. Plates were then re-sealed and put back in growth chambers, with settings mentioned above. After 28 more days in culture, plants were phenotyped by measuring their shoot fresh weight.

### Biolog Plate Experiment

Mycelia of individual *Plectosphaerella* isolates (n = 72) were harvested using sterile tips and transferred into 1 mL of 10 mM MgCl₂ containing one stainless-steel bead (Ø = 2.85–3.45 mm; Carl Roth) and ten smaller beads (Ø = 0.75–1 mm; Carl Roth). Samples were homogenized using a paint shaker (SK450, Fast & Fluid Management, Sassenheim, Netherlands) for 10 minutes. The resulting fungal suspensions were diluted to a final biomass concentration of 0.2 mg/mL in a 1:1 mixture of Biolog FF and IFO inoculating fluids (Biolog Inc.), supplemented with 1% Dye Mix A (Biolog Inc.) to enable colorimetric detection. For each isolate, 100 µL of the inoculum was loaded into Biolog PM1 and PM2A MicroPlates™, which together contain 190 distinct carbon sources. Plates were incubated at 25 degrees without shaking and respiration was monitored every 20 minutes for 5 days using the OmniLog® instrument (Biolog Inc.), which records OmniLog Respiration Units (ORU). The area under the curve (AUC) of ORU values was calculated for each well over the 5-day period and log-transformed (log AUC) for normalization prior to data visualization and statistical analysis.

### Genome sequencing

Two strains in our collection (P16 and P117) were previously sequenced with the PacBio technology and assembled^8^. Their genomes made available on database GenBank: bioprojects PRJNA371204 and PRJNA570880. While we used previously published genome assembly JAGPXD000000000 for P16, we used an improved version of JAGSXJ000000000 for P117, which was downloaded from the Mycocosm portal^85^ with the identifier Plecucu2. For P30, we used a genome assembled from Illumina short reads made available on the Mycocosm portal with identifier Plecu1. Other genomes used in this study were newly sequenced with PacBio sequencing. Individual strains were revived on Potato Growth Agar medium (Roth, ref. CP74.1), and cultured for 7 days. Six strains (P43, P143, P340, P421, P455, P612) were sequenced with PacBio CLR technology. Their gDNA was extracted according to a previously introduced cetyltrimethylammonium bromide (CTAB) protocol^32^. RNA of these strains was sequenced with Illumina short reads to support gene prediction in the assemblies. All the other strains (n = 63) were sequenced with PacBio CCS technology (HiFi reads). Their gDNA was extracted according to a previously introduced CTAB protocol^32^, without the described purification step of the DNA extracts. Native fungal DNA was found in many cases to be recalcitrant to sequencing. Therefore, PacBio barcoded ultra-low libraries were prepared according to the protocol “Procedure Checklist - Preparing HiFi SMRTbell® Libraries from Ultra-Low DNA Input” starting with DNA fragmentation using g-Tubes (Covaris). Typically, five libraries were pooled and then sequenced on a Sequel IIe SMRT cell with 8 million ZMWs with Binding kit 2.0 or Binding kit 2.2 and the Sequel II Sequencing Kit 2.0 for 30 h. If necessary gDNA or final libraries were additionally size-selected on a Blue Pippin (Sage Sciences) device to remove smaller DNA fragments. After sequencing, the HiFi data was demultiplexed with SMRTlink10.0, and adapter sequences were trimmed away with cutadapt v3.5^86^.

### Genome assembly and gene annotation

Genomes were assembled and polished from PacBio reads using Flye v2.9-b1768^87^ with parameter *–genome-size 40m*, decided based on the size of previously sequenced *P. cucumerina* genomes^8, 30^. Parameter –pacbio-hifi was used for genomes sequenced with PacBio CCS and –pacbio-raw was used for genomes sequenced with PacBio CLR. Gene prediction was carried out in all the genome assemblies except for P16 and P117, for which we used the gene predictions from the JGI annotation pipeline^85^. We used FGENESH v8.0.0^88^ with similarity matrix *Torrubiella hemipterigena* and parameters *-skip bad prom-skip bad term*. CAZyme-encoding genes were annotated using dbcan v4.0.0^89^. Candidate effector-encoding genes were annotated by first predicting secreted proteins with SignalP 6.0^90^, then by using EffectorP v3.0^91^ (in fungal mode) on predicted secreted proteins. To annotate protease-enconding genes, the annotation tool emapper v2.1.5^92^ based on eggNOG orthology data^93^ was run using DIAMOND as a similarity search algorithm^94^, then we considered as proteases all the proteins linked by emapper to a PFAM identifier associated to a MEROPS family^95^. Transposons were annotated in genome assemblies with reasonaTE v1.0^96^ with parameters *- mode annotate-tool all*.

### Comparative genomics

Orthology prediction was performed on the total set of proteins predicted in our data set using OrthoFinder v2.5.4^97^ with parameter -S blast. Orthogroups were classified into intra-species conservation levels. Those represented in 100% *P. cucumerina* genomes were called *core*. The ones represented in at least 75% *P. cucumerina* genomes were considered *broadly conserved*. The ones represented in less than 20% *P. cucumerina* genomes were considered to have low conservation. We considered as *subspecies-associated* the orthogroups represented in more than 20% of one *P. cucumerina* subspecies, while absent from the other. Remaining orthogroups were considered as *sparsely distributed*. To test for overrepresentation of these conservation levels among orthogroups encoding CAZymes, effectors and proteases, Fisher’s exact tests were performed using the function stats.fisher_exact(alternative=’two-sided’) from Scipy^80^. Principal components analysis (PCA) revealing similarities of CAZyme, protease and effector catalogues were performed by counting the number of annotated genes per family (CAZy and MEROPS families for CAZymes and proteases respectively, and orthogroups for effectors), then calculating pairwise Jaccard distances between strains. Jaccard dissimilarity matrices were calculated with R package Vegan v2.6-4 (https://github.com/jarioksa/vegan), using function vegdist(method=”jaccard”). PCA plots were then reconstructed from these matrices using function prcomp in R. PERMANOVA tests were computed to test for effects of strain phylogeny and the host of isolation on CAZyme, effector and protease catalogues, using the function adonis2(JaccardMatrix∼PhylogeneticGroup+HostOfIsolation, permutations = 9999). Following PERMANOVA, given the significant effect of *Plectosphaerella* phylogenetic groups on genomic compositions in gene repertoires, pairwise comparisons aiming to detect significant differences between phylogenetic groups were computed with the function pairwise.perm.manova() from R package RVAideMemoire v 0.9-83-12, with default Benjamini-Hochberg correction applied on *P* values.

### Phylogeny reconstruction

We used four different methods to reconstruct the phylogeny of our *Plectosphaerella* collection, based on genome sequences. First two trees were built from 5,466 single copy orthologs, identified in our data set by OrthoFinder^97^. Sequences of these 5,466 gene families were aligned independently using MAFFT v7.407 with parameter –auto^98^, then trimmed with Trimal v1.4.rev22 using default settings^99^. These trimmed alignments were used as input for IQ-TREE v2.1.1^100^. The algorithm ModelFinder^101^ implemented in IQ-TREE identified maximum-likelihood model JTT+F+I+G4 to be the most adapted one considering the input data. We therefore reconstructed phylogenies with IQ-TREE and RAxML-NG v1.1.0^102^ using this model. While prior alignment concatenation using AMAS^103^ was necessary in the case of RAxML-NG, the IQ-TREE software carries out concatenation before phylogeny reconstruction. Additionally, we reconstructed phylogenies with two different coalescent approaches. First, we used method STAG^97^, as implemented in OrthoFinder. STAG built individual gene family trees (n=13,694), then reconstructed a coalescent species tree from them. Finally, we used Astral v5.7.1^104^ with default parameters, to reconstruct a coalescent species tree from all the single copy ortholog gene trees generated by OrthoFinder (n = 5,466). The phylogenetic tree used as a reference in this study and presented on multiple figures is the one generated by IQ-TREE.

### Multi-host re-colonization experiments with *P. cucumerina* 0016

Arabidopsis and tomato seeds were sterilized for 15 min in 70% ethanol, then 5 min in 8% sodium hypochlorite. Subsequently, the seeds were rinsed five times with sterile double-distilled water and once with a 10 mM Magnesium Chloride (MgCl_2_) solution. They were then stratified in darkness at 4°C for a period of 2 to 3 days. Barley seeds were sterilized for 1 hour in 6% sodium hypochlorite and then five washes in sterile double-distilled water. Before inoculation, Arabidopsis and tomato seedlings were grown for 14 days and barley seedlings were grown for 7 days in ½ Murashige and Skoog (MS) media (without carbon source, pH 5.7). The conditions in the growth chamber are 21°C, for 10h with light (intensity 4) at 19°C and 14h in the dark in growth chambers. Inoculation in the roots with *P. cucumerina* 0016 was carried out by harvesting the spores from mycelium grown for 8 days on Potato extract Glucose Agar medium (PGA) using sterile double-distilled water. Harvested spores were filtered through Miracloth (Merck, 475855-1R) centrifuged for 5 min at 600 x g (Eppendorf-centrifuge 5810R) and then pellets were resuspended in 1 mL of sterile double-distilled water. Spores were counted using Hemocytometer and the concentration was adjusted to 2.5 x 10^6 spores/mL. These spores were inoculated onto the roots of Arabidopsis, tomato, and barley seedlings. After 30 min of incubation, the seedlings were transferred to sterile Magenta boxes containing 40 g vermiculite with 95 mL of ½ MS media (without carbon source, pH 5.7). After transferring four plants into this “vermiculite system”, the boxes were sealed using Micropore by placing an empty magenta box upside down on top. Plants (Arabidopsis, tomato, and barley) were grown for 21 days post inoculation (dpi) in the same growth conditions as described above. As a control condition for RNA-seq experiment, sterile Nucleopore Track-Etched polyester membranes (“FungusOnMembrane”) were deposited on “Vermiculite System”, then 10 μL drops of fungal spores (2.5 x 10^6 spores/mL of *P. cucumerina* 0016 in sterile double-distilled water) were placed on each one. The membranes were collected and processed as the root samples of our test condition. After 21 dpi, the roots were harvested and flash-frozen (if they were for *in planta* RNA seq experiment), shoot fresh weight (SFW) was measured for each plant. For analysis, the plant performance index was calculated as the product of shoot fresh weight (SFW) and survival rate of the plants^8^. In further correlation analyses, we used log transformed PPI normalized to mock controls using the Hedges’ g method. The harvested roots were stored at -80°C until further use.

### High-throughput DNA extraction and quantification of fungal load at the roots

Total DNA was isolated from Arabidopsis, tomato and barley frozen roots samples (one root per tube) using a modified high-throughput version of the FastDNA SPIN kit for Soil (MP Biomedicals)^105^. Root samples were homogenized in a Precellys 24 TissueLyser (Bertin Technologies) for 2 x 30 s at 6,200 rpm with 15 s intervals. 450 µL of sodium phosphate buffer (MP Biomedicals) and 50 µL of MT buffer (MP Biomedicals) was added and then homogenized as described before. Subsequently, the centrifugation was performed for 15 min at 16,000 x g (Eppendorf-centrifuge 5424R) and 150 µL of the supernatant were transferred to a 96-well Acroprep Advance filter plate (with 0.2 μm Supor filter; Pall). Once full, the filter plate was positioned on a PCR plate filled with 50 µL Binding Matrix (MP Biomedicals) per well and centrifuged for 15 min to remove residual soil particles. This and all subsequent centrifugation steps were carried out in a swing out centrifuge at 200 x g (Eppendorf-centrifuge 5424R). The PCR plate was sealed and shaken for 3 min to allow binding of the DNA to the Binding Matrix. The suspension was pipetted onto a second filter plate of the same kind, positioned on a collection plate, and centrifuged for 15 min. The flowthrough was discarded. Then, 200 µL SEWS-M washing buffer (MP Biomedicals) were pipetted into each well of the filter plate and centrifuged for 5 min. This washing step was carried out a second time. The flowthrough was discarded and followed by centrifugation for 5 min to remove residual SEWS-M buffer. Finally, 30 µL nuclease-free water were added to each well and left to incubate at room temperature for 3 min. Subsequent centrifugation for 5 min eluted the DNA into a clean PCR plate. Fungal colonization of these root samples was then measured by quantitative PCR (qPCR) without prior adjustment of DNA concentrations. For each Arabidopsis root sample, two reactions were conducted with one pair of primers ITS1F (5’-CTTGGTCATTTAGAGGAAGTAA-3’) and ITS2 (5’-GCTGCGTTCTTCATCGATGC-3’) which target the fungal ITS1 sequence, and two pair of primers specific to plant samples: UBQ10F (5’-TGTTTCCGTTCCTGTTATCT-3’) and UBQ10R (5-ATGTTCAAGCCATCCTTAGA-3) that target the *Ubiquitin10* Arabidopsis gene, nAP256F (5’-CTACCACTGGTCACTTGATC-3’) nAP257R (5’-CCAGGCATACTTGAATGACC-3’) that target the *EF1-alpha* tomato gene, UBQF (5’-ACCCTCGCCGACTACAACAT-3’) and UBQR (5-CAGTAGTGGCGGTCGAAGTG-3’) that target the *Ubiquitin* barley gene^8, 106^. Each reaction was performed by mixing 5 μL of iQ™ SYBR Green Supermix with 0.3 μL of 10 μM forward primer, 0.3 μL of 10 μM reverse primer, 3.4 µL of sterile double-distilled water and 1 μL of template DNA. A BioRad CFX Connect Real-Time system was used with the following program: 3 min of denaturation at 95°C, followed by 39 cycles of 15 s at 95°C, 30 s at 60°C and 30 s at 72°C. We then calculated a single colonization index for each sample using the following formula: Index = 2−(Cq(ITS1)/Cq(UBQ10))^8^.

### Confocal microscopy of root colonization by fungi

Roots of Arabidopsis (5 weeks old), tomato (5 weeks old), and barley (4 weeks old) grown for 21 days in mono-association with *P. cucumerina* 0016 were harvested and conserved in 70% ethanol. During the process, the roots were incubated in 20% of Potassium Hydroxide (KOH) at room temperature with orbital shaking at 60 rpm overnight. At the end of the incubation, the KOH was removed and the roots were rinsed twice with 1x Phosphate-Buffered Saline (PBS) at pH 7.4. They were then double-stained with 0.1% Calcofluor white (Sigma, cat: 18909) and 25 µg/ mL (for Arabidopsis roots)-50 µg/mL (for tomato and barley roots) of Wheat Germ Agglutinin conjugated to fluorophore Biotium CF633 (WGA-CF633) buffered at pH 7.4 in PBS. After 30 mins of incubation, samples were then washed thrice in PBS and imaged with a Confocal LSM880 from Zeiss and the associated software ZEN v2.3 SP1.

### *In planta* fungal transcriptome analysis

PolyA enrichment was carried out on the RNA extracts, then RNAseq library was prepared with the NEBNext Ultra™ II Directional RNA Library Prep Kit for Illumina (New England Biolabs). Sequencing was then performed in single read mode on a HiSeq 3000 system. RNA-seq reads from Agar System were trimmed using Trimmomatic^107^ v0.38 and parameters TRAILING:20 AVGQUAL:20 HEADCROP: 10 MINLEN:100. For RNA-seq reads from Vermiculite System, trim-galore v0.6.5 pipeline was used^108^. Subsequently, these reads were employed for the quantification of transcripts in each sample. A unified transcriptome was constructed by concatenating the transcriptomes of plants and *P. cucumerina* (Plecuc1_GeneCatalog_CDS_20171208 from JGI Mycocosm^8, 85^) for various hosts. The indexing for Arabidopsis was based on TAIR11^109^ from JGI-Phytozome^110^, for tomato was based on TMCSv1.2.1^111^, for barley was based on Hvulgare_462_r1^112^. The creation of an index for the concatenated transcriptome was accomplished using Salmon v1.1.0^113^. The quantification of transcripts was performed with Salmon quant to determine the read counts for each sample. These datasets were then imported into R utilizing the tximport v1.12.3^114^. Bioconductor package to compute the Transcript Per Million (TPM) and Reads Per Kilobase per Million mapped reads (RPKM) values. Package *DESeq2* v1.24.0^115^ was used for quantification of differential expression of genes (DEGs) and log_2_ fold-change (Log2FC) was calculated using “FungusOnMembrane” as control group. Log2FC values were corrected by shrinkage with the algorithm in *apeglm* v1.6.0 package^116^. Fungal DEGs obtained on infecting different host conditions were used for further analysis. Subsequently, a matrix was created by incorporating the log_10_ (counts per million, CPM) values for individual samples (columns) along with exclusively the fungal DEGs (rows). Computation of log_10_ CPM values was based on the normalized read counts.

### Statistical analysis of differentially expressed genes

Using the computed CPM matrix, the Pearson correlation among samples was determined and was used as the basis for multi-dimensional scaling (MDS) analysis. MDS was carried out utilizing the distance matrix, and pairwise statistical analyses were conducted using Dim1 and Dim2 for each sample group to assess the significance of impact across conditions, employing the ANOVA-TukeyHSD function in R. This matrix was also employed for clustering the DEGs using *k*-means clustering algorithm and the optimum *k* (ranging from 1 to 100) was determined using the Akaike Information Criterion (AIC) and Bayesian Information Criterion (BIC). A heatmap was generated using the Log2FC to demonstrate expression patterns in comparison to the control. Visualization of DEG within the groups was illustrated through VennDiagram^117^ and heatmap^118^ separately. Gene annotations were assigned by matching the respective gene/transcript/protein IDs to available annotations in JGI Plecuc1 (*Plectosphaerella cucumerina* RP01 v1.0^8^) that includes Gene Ontology (GO), INTERPRO, EffectorP^91^, and CAZyme^119^ terms associated with the fungal genes. For each functional term, gene set enrichment analysis was performed using Fisher’s Exact test. The *P* values were adjusted for each condition using the Benjamini-Hochberg method.

### Construction of CAZyme network plots targeting predicted Cell Wall Substrates

CAZyme database was utilized for CAZyme enrichment analysis and visualization. Substrates were categorized manually based on the class of Cell Wall (CW) (string parsing *“pectin|cellulose|lignin|arabinan|galactan|glycan|glucan|xylan”* using *tidyverse*) components, and redundant substrates were grouped together to simplify visualization. A network graph was created using an adjacency matrix consisting of CAZyme families (rows) and possible substrate families (columns) (CAZyDB vCAZyDB.09242021 from dbCAN2 repository)^47^. A value of 1 was assigned to CAZyme families targeting potential substrates, with the rest being designated as 0. Using this matrix network map was constructed, where the edges indicated the number of enzymes (circle nodes) targeting potential substrates (square nodes). *P* values derived from the enrichment analysis of the corresponding CAZyme families were employed to represent the nodes in the network graphs, mean Log2FC of the CAZymes was employed to illustrate the color of the edges. By integrating the network structure and the significantly enriched CAZymes, the activation of CW components under specific conditions was visualized using the igraph^120^ package in R (v3.6.1 & v4.3.1).

### Monosaccharide compositional analysis and crystalline cellulose measurement

Cellulose content and matrix polysaccharide monosaccharide composition of the samples were analysed as per previously described method^121^. In short, Alcohol insoluble residue (AIR) was prepared from freeze-dried root tissues and split into two samples. One half of the samples were treated with a weak acid (4% sulfuric acid) to release matrix polysaccharide-derived sugars, while the other half of the samples were treated first with a strong acid (72% sulfuric acid) to swell cellulose. Then the sulfuric acid concentration was diluted to 4% to yield monosaccharides both derived from cellulose and the matrix polymers. Subtraction of the two values allows for the quantification of crystalline cellulose. Monosaccharides of all fractions were quantified using a Professional IC Vario high-performance anion-exchange chromatography system 1068 (Metrohm, Herisau, Switzerland) equipped with a CarboPac PA20 column (Thermo Fisher Scientific, Waltham, MA, USA) and an amperometric detector (Metrohm). The used sodium hydroxide gradient is specified as described^121^. Lignin content of the AIR samples was determined as previously described^122^.

### GH64 occurrences in the fungal kingdom

The CAZYme family GH64 was procured from the JGI Mycocosm^85^ and from the CAZyme database^119^ respectively. These datasets were merged to preserve the taxonomic categorization for the fungal kingdom that were present in the CAZyme database based on the portal ID (genomic identifiers). The number of GH64 occurrences was ascertained by cross-referencing the merged dataset. Finally, the FunGuild database was integrated into the concatenated dataset through species or genus identifiers. Collectively, three tables were merged and summarised using a phylogenetic tree of the fungal kingdom (constructed from the *taxonomizr* R package^123^). The count data for the number of GH64 in genomes (from JGI Mycocosm^85^) and the ecological information (FunGuild^124^) were incorporated into the phylogenetic tree utilizing iTOL v6.0^125^. Due to redundancy in the annotation of ecological niches (GUILD category in the FunGuild repository^124^), we classified those annotations that contain the string “endophyte” as endophyte while others were classified as “non-endophyte.” Statistical analysis for Phylum was executed employing the pairwise Wilcoxon test with Bonferroni *P* value adjustment, and for enrichment analysis within Ascomycota members (at the class level) and within the ecological group, Fisher’s exact test was performed with *P* value adjustment using Bonferroni correction.

### Structural analysis of GH64

The structure of the GH64 protein from *P. cucumerina* 0016 was predicted with AlphaFold3, after signal peptide removal with SignalP v6.0^90^. The best predicted model was then used as query for Foldseek^50^ search (online, accessed December 2024), looking for structural similarity with experimentally resolved protein structures registered in the PDB database. Structure alignment between the GH64 enzyme of P16 and the best hit (SpGlu64A, protein from *Streptomyces* pratensis^51^, PDB code: 7VTK) was performed using TM-align^126^, as implemented in iCn3d^127^.

### CRISPR/Cas9-mediated genome editing in *P. cucumerina* 0016

#### Design of synthetic guide RNA (sgRNA)

The structure of the specific sgRNA oligo is T7 promoter-Start of transcription-Protospacer-Constant part of sgRNA. AAGCTAATACGACTCACTATAG(G)NNNNNNNNNNNNNNNNNNNNGTTTTAGAGCTAGAA ATAGCAAG. sgRNAs (**Supplementary Table 5**) were designed with the sgRNA designer tools CRISPick (https://portals.broadinstitute.org/gppx/crispick/public) and CCTop^128^. An sgRNA with a single, unique genomic hit to the target locus was identified using CCTop, and its on-target score was retrieved from CRISPick.

#### Synthesis of sgRNA

For the preparation of gRNA synthesis, specific and constant gRNA oligos were mixed by combining 1 µL of the specific protospacer oligo (100 pmol stock), 1 µL of the constant gRNA oligo (100 pmol stock), and 8 µL of H2O, totaling 10 µL. 95°C for 5 min, cooling from 95°C to 85°C at 2°C/sec, and further cooling from 85°C to 25°C at -0.1°C/sec, with a total time of approximately 15 min. A T4 DNA mix was prepared to fill in overhangs, consisting of 2.5 µL dNTPs (10 mM stock), 2 µL NEB buffer 2.1 (10x buffer stock), 5 µL H2O, and 0.5 µL T4 DNA polymerase (NEB, M0203L), totaling 10 µL. This T4 DNA mix was added to the annealed oligos, and the mixture was incubated at 12°C for 20 min. The product was purified using a PCR column (Macherey-Nagel NucleoSpin Gel and PCR Clean-up Kit) and eluted with 30 µL H2O. The concentration was measured with a Nanodrop, ensuring it was around 130 ng/µL, and the band size was verified on a 3% agarose gel, expecting a band of approximately 120 bp. For RNA synthesis, the HiScribe T7 High Yield RNA Synthesis Kit (Neb, E2040S) was used, following the protocol for small RNAs. The reaction mixture included 6 µL NTPs (25 mM stock), 1.5 µL T7 Buffer (10x buffer stock), 1.5 µL T7 Polymerase, 11 µL Template (1-2 µg DNA) and, totaling 20 µL (overnight incubation at 37°C). After that, 14 µL of nuclease free H2O, 4 µL of DNase 1 buffer (10x), and 2 µL of DNase 1 were added, and the setup was incubated for 30 min at 37°C.

#### Purification of sgRNA

For sgRNA purification, RNA Clean & Concentrator 25 kit (Zymo Research, Catalog No R1017 & R1018) was used. The eluted samples were immediately stored at -80°C, with a concentration greater than 1 µg/µL. The integrity of the gRNA was checked on 3-4% agarose gel electrophoresis. If a single band was observed, it was suitable for further steps. If a smear was present, the sgRNA synthesis needed to be repeated. If a double band was observed, double the amount of sgRNA than recommended was used for transformation.

#### Design of the Donor DNA

Transgenes (donor DNAs) were designed to include a homology arm on the 5’ site of the Cas9 cutting site (60 bp) (**Supplementary Table 5**), the promoter Pcpc1, the phleomycin resistance gene *BleoR*^129^, the transcription terminator of tryptophan synthase (TtrpC), and a homology arm on the 3’ site of the Cas9 cutting site (60 bp). This construct was ordered from the company Biocat (https://www.biocat.com/gensynthese) in plasmid pUC57.

#### Amplification of Donor DNA using PCR

Once the construct was received, donor DNA amplification was performed using PCR. Prior to this, primers (**Supplementary Table 5**; P5-P16 used in pairs) were designed to amplify the donor DNA (*Bleo* cassette with the flanking region, as described) from the vector. The reaction conditions included 5 µL of Buffer (5X), 0.4 µL of dNTPs (25 mM), 1 µL of DNA Template, 1 µL of Forward Primer (100 µM), 1 µL of Reverse Primer (100 µM), 1 µL of Phusion DNA polymerase, and 41.6 µL of Sterile water. The PCR conditions were set as follows: 98°C for 30 s, followed by 35 cycles of denaturation at 98°C for 8 s, annealing at required temperature for 30 s, and extension at 72°C for 45 s, with a final extension at 72°C for 10 min. A 1% agarose gel electrophoresis with a 1 kb ladder (GeneRuler, Thermo Fisher Scientific) was run to check the integrity of the DNA, ensuring a single clean band was obtained. The DNA band was cut from the gel, and gel purification was performed using the Macherey Nagel kit. The concentration was checked using a Nanodrop, ensuring it was greater than 200 ng/µL.

#### Protoplast isolation

Spores at a concentration of 1.5 x 10^7 cells/ mL were inoculated into 100 mL of PGB medium. The flask was incubated at 25°C with orbital shaking at 180 rpm for 20 h. Next day, the mycelia were collected by passing the entire culture through a 40 µm filter, and the residues were washed vigorously using 20 ml of 0.7M NaCl to completely remove the spores from the mycelia^130^. Mycelia (350 mg) were transferred to 10 mL of enzyme solution (20 mg driselase in 10 mL of 0.7 M Nacl) and the mixture was incubated in the enzyme solution for 2 h at 28°C with orbital agitation at 60 rpm. After incubation, the solution was filtered (40 µm filter). The filtrate was centrifuged at 250 x g for 5 min, with acceleration and deceleration adjusted to 3 in cg (Eppendorf-centrifuge 5810R). The pellet was resuspended in 1 ml of STC (Sucrose 20% (w/v), TrisHcl (1M, pH 8)-10 mM, Cacl2 (1M)-50 mM) buffer. The protoplasts were counted using a hemocytometer, and the volume was adjusted to 5 x 10^6 cells/mL. *PEG mediated transformation:* For the PEG-mediated transformation, prebound Cas9-gRNA complex (RNP-Ribonucleoprotein complex)^131^ was prepared by mixing Cas9 and sgRNA as follows: 0.5 μL of 10X NEBuffer r3.1, 1 μL of 20 μM EnGen Spy Cas9 HF (NEB, M0667M), and 1.5 μL of sgRNA 2 (approximately 1.5 µg, with the desired amount being 20-40 pmol, or 0.8-1.6 μg for 121 nucleotides). This whole mixture was incubated at 25°C for 30 min and further kept on ice until use. For protoplast transformation, 200 µL volume of protoplasts were taken in a 50 mL Falcon tube and kept on ice. The pre-bound Cas9-gRNA complex (∼20 pmol) and donor DNA (x µL containing 2.5 µg of DNA) were added to the same tube and gently mixed, the total volume should be as low as possible. The tube with mixture was incubated on ice for 30 min. Then, 1.5 mL of PEG-STC (6 g of PEG 8000 in 14 mL of STC buffer) was added to the tube, which was rotated horizontally to softly mix the reagent. This was followed by incubation at room temperature for 15 min. Subsequently, 5 mL of CM media (containing 6 g of yeast extract, 6 g casein acid hydrolysate, and 10 g sucrose in 1 L H_2_0) was added and mixed well by horizontal rotation. The tube was incubated in the dark at 25°C for 18 h before being centrifuged at 3,000 x g for 5 min. The pelleted fungi were visible on the side of the tube, and the supernatant was removed. Regenerated protoplasts were resuspended together in 100 μL of sterile water the entire solution was plated on square plates containing PGA with 200 µg/mL *Bleo* antibiotic and incubated in a 23°C incubator. After 2 days, these colonies were transferred to new plates (PGA with *Bleo*), marked with numbers, and allowed to grow for 4-5 days before proceeding with screening the mutants. For control, untransformed protoplasts were plated on PGA and used for screening steps to maintain uniformity for the wild type samples and transformed colonies, especially in the colony PCR step. To screen and to verify the integration of *BleoR* gene cassette at the right site, we used external primers that anneal to the genomic sequences immediately outside the 5′ and 3′ homology arms (**Supplementary Table 5)**. For colony PCR, the Phire Plant Direct PCR Master Mix was used (Thermofisher, F170S). Mycelium was picked with a tip and it was transferred into 30 µL of the dilution buffer. This was incubated at 95°C for 15 min and then overnight at 4°C. The next day, the template was ready. The reaction mixture as follows: 10 µL of Master Mix, 1 µL of 10 µM forward primer, 1 µL of 10 µM reverse primer, 0.5 µL of Template, and 7.5 µL of nuclease free water. Once the PCR was done, agarose gel electrophoresis was run to check the results and once a positive colony was identified, DNA extraction was performed using the Qiagen DNeasy Plant Mini kit, and PCR was conducted again with Phusion polymerase to double-check the results.

### *In vitro* growth assay

To assess the growth rates of mutant *P. cucumerina* 0016 strains relative to the wild type, an *in vitro* growth assay was conducted. Spores were initially collected from mycelia grown on PGA. Using a hemocytometer, the spores were counted, and their concentration was adjusted to 2.5 x 10^6 spores/mL. These spores were then inoculated into 30 mL of PGB and incubated at 23°C with orbital shaking at 180 rpm for 24 hours. After incubation, a 1 mL sample was taken to measure the optical density (OD) at 600 nm with a spectrophotometer. The remaining culture was used to harvest and recount spores using a hemocytometer. This experiment included three biological replicates. *In vitro* growth dynamics of *P. cucumerina* 0016 wild-type (WT) and Δgh64 mutant strains in PGB medium were performed in 96 well-plates and measured using spectrophotometer. For WT and Δgh64 conditions, 180 µl PGB was dispensed per well whereas for mock 200 µl PGB was dispensed per well. The outer wells were filled with water to minimize edge effects. Mycelia were harvested from PGA and then they were washed with water to remove residual agar. Further, they were resuspended in water with the concentration adjusted to 2 mg/ml. Later, they were homogenized using a Precellys 24 TissueLyser (Bertin Technologies) at 6,200 rpm for 2 × 30 s with a 15 s interval. Subsequently, 20 µl of mycelial suspension was added to wells containing 180 µl PGB. Each condition was represented by six wells per experiment and three independent biological experiments were conducted. Optical density at 600 nm (OD600) was measured at 0, 24, 48, 72, and 96 h using microplate reader (Tecan, Infinite M Nano). For the agar plate assay, 20 µl of mycelial suspension (2 mg/ml) was placed at the centre of PGA plates for each condition, and mycelial growth were recorded after 10 days. For both the experiment, the samples were incubated at 23°C.

### Experiments with C. incanum

#### Targeted gene replacement

*Agrobacterium*-mediated transformation was used based on Hiruma et al., 2023^60^. Briefly, the upstream and downstream (∼1500 bp each) regions surrounding the *GH64* gene in *Colletotrichum incanum* MAFF 238704 (*Ci*) were amplified by polymerase chain reaction (PCR) using the PrimeSTAR HS DNA polymerase (TaKaRa). PCR products were electrophoresed on a 1% (w/v) agarose gel, then DNA was extracted from the cut gel using the FastGene™ Gel/PCR Extraction Kit (NIPPON Genetics). The purified upstream and downstream regions were mixed with SalI-treated pBIG4MRHrev using the In-Fusion® HD Cloning Kit (TaKaRa). The generated plasmid was subsequently introduced to *Agrobacterium tumefaciens* C58C1 by electroporation (1800 V, 5 ms) using the Eppendorf Eporator. *Ci* spores were prepared by culturing agar plugs from Mathur’s agar (3% w/v) plates into Mathur’s agar flasks, incubating for 2 weeks, then harvested by flooding the flask with GI broth supplemented with 200 μM acetosyringone before washing twice with GI broth supplemented with acetosyringone. Transformed *Agrobacterium* (OD_600_ = 0.4) was mixed 1:1 with Ci spores (10^7^ spores/mL) on sterile paper filter discs on GI agar (1.5% w/v) supplemented with 200 μM acetosyringone. Two days after transformation, filter paper discs were flipped onto Mathur’s agar plates supplemented with hygromycin (200 μg/mL), cefotaxime (100 μg/mL), and spectinomycin (100 μg/mL) and incubated for 2 days at room temperature (∼25°C) before filter papers were removed. Colonies were allowed to merge, then were re-streaked onto fresh Mathur’s plates supplemented with hygromycin, cefotaxime, and spectinomycin (same concentrations). Colony PCR was performed to validate the success of the gene replacement using primers listed in **Supplementary Table 5**.

#### Seed inoculation

Agar plugs from fungal cultures streaked on Mathur’s agar plates were transferred to fresh Mathur’s agar plates and incubated at 25°C for 7 days under continuous light (∼10 μmol m^-2^ s^-1^). *A. thaliana* Col-0 or *Nicotiana benthamiana* seeds were surface sterilized with 70% ethanol for 30 seconds twice, followed by 6% sodium hypochlorite with 0.01% Triton X for 5 minutes. After being washed 5 times with sterilized double-distilled water (DW), seeds were sowed on half-strength Murashige and Skoog (MS) agar (1.3% w/v) plates without sucrose. Spores from Mathur’s plates were collected by flooding the fungal culture plate with 5 mL DW, washed once in DW, then counted by hemocytometry in a light microscope. Spores were diluted to 5,000 spores/mL prior to inoculating 3 μL of the spore suspension 3 cm beneath seeds. Plates were incubated vertically in a plant growth chamber at 22°C under fluorescent light (∼100 μmol m^-2^ s^-1^) with a 16-hour photoperiod at ∼40% humidity. Seedlings were harvested at 24 dpi for shoot fresh weight measurement. Roots were sampled, flash frozen in liquid nitrogen, and stored at -80°C until RNA was extracted using the Nucleospin^®^ RNA kit (Macherey-Nagel) according to the manufacturer’s instructions. RNA was immediately used for cDNA synthesis, then stored at -80°C.

#### Quantitative real-time polymerase chain reaction (qRT-PCR)

cDNA was synthesized from ∼100 ng total RNA using the PrimeScript™ RT Master Mix (TaKaRa) in a volume of 10 µL following manufacturer’s instructions. qRT-PCR was performed using the THUNDERBIRD^®^ Next SYBR qPCR Mix (TOYOBO) with 250 nM primers using CFX Opus 384 Real-Time PCR System (BIORAD) in a volume of 10 µL for 48 cycles. Primers are listed in **Supplementary Table 5**.

### Statistics

For the graphs and statistical analysis, the R programming environment (R version 4.4.0) was used for data analysis and visualization (https://www.r-project.org/). The normality of the data was assessed with the Shapiro–Wilk test using stats package (V4.4.0) and homogeneity of variances with Levene’s test using car package (V3.1-2). The two-sided T-tests and two-sided non-parametric Wilcox rank-sum test were performed using t.test and wilcox.test from the stats package (V4.4.0) in R. ANOVA-TukeyHSD tests were conducted using emmeans package (V1.10.3). Non-parametric Kruskal-Wallis tests were carried out with stats package (V4.4.0), and Dunn’s tests were performed using FSA (V0.9.5) and rcompanion packages (V2.4.36). Reported *P* values were Bonferroni adjusted using stats package (V4.4.0).

## Data availability

RNASeq raw data are available at the European Nucleotide Archive (ENA) under the study ID PRJEB89666. The raw genomic data of the *Plectosphaerella* isolates have been deposited in the ENA database under the project IDs PRJEB61839 (P143), PRJEB50298 (P340, P421, P455) and PRJEB90544 (all other isolates). See **Supplementary Table 3** for individual ENA sample IDs.

## Code availability

Scripts and raw data used to generate figures are available at https://github.com/fantin-mesny/Scripts_comparison_of_Plectosphaerella_strains for WGS, https://github.com/arpankbasak/Plecuc1_Repo. For RNAseq, and https://github.com/RamSevakRK/Genetic-factors-driving-multi-host-infection-in-a-core-member-of-the-root-mycobiota for all the other experiments.

## Acknowledgements

This work was supported by funds to S.H. from European Research Council consolidator grants MICRORULES (758003) and MICROBIOSIS (101089198), the Cluster of Excellence on Plant Sciences (CEPLAS) under Germany’s Excellence Strategy-EXC 2048/1-Project ID: 390686111, and the Priority Programme: Deconstruction and Reconstruction of the Plant Microbiota (SPP DECRyPT 2125; HA 8169/2-2), both funded by the Deutsche Forschungsgemeinschaft (DFG, German Research Foundation). A.K.B. was funded by the Polish National Agency for Academic Exchange NAWA (BPN/BEK/2023/1/00315). J.N. obtained funding from a JST SPRING grant (JPMJSP2108) and K.H. from a JST grant (JPMJAN23D4). J.G.M-V. is funded by the Beatriz Galindo program (BG23/00164) of the Spanish Ministry of Science, Innovation and Universities. We would like to thank Khushi Joshi (MPIPZ), Thorsten Thiergart (MPIPZ), and Katharina Grosche (Heinrich-Heine University) for the excellent technical assistance and Bart Thomma (University of Cologne) for providing protocols for fungal genome editing.

## Author contributions

Conceptualization: S.H., F.M., and R-S.R-K. Supervision: S.H. Methodology and Investigation: R-S.R-K., A.B.K, F.M., S.H., J.N., G.C., F.E., L.R., S.C.A., M.P., and K.H. Analysis Support: T.L. and T.T. Sequencing Support: B.H. Contribution to biological material: P.W.C., J.G.M-V., H.S., M.R., H.R., A.M.F., S.S., A.M., I.B., S.D., W.H.E., J.H., J.S.K., J.M.N., and C.M.S. Visualization: R-S.R-K., A.B.K, F.M. and S.H. Writing original draft: S.H and R-S.R-K.

## Supplementary Figures

**Supplementary Fig. 1.**
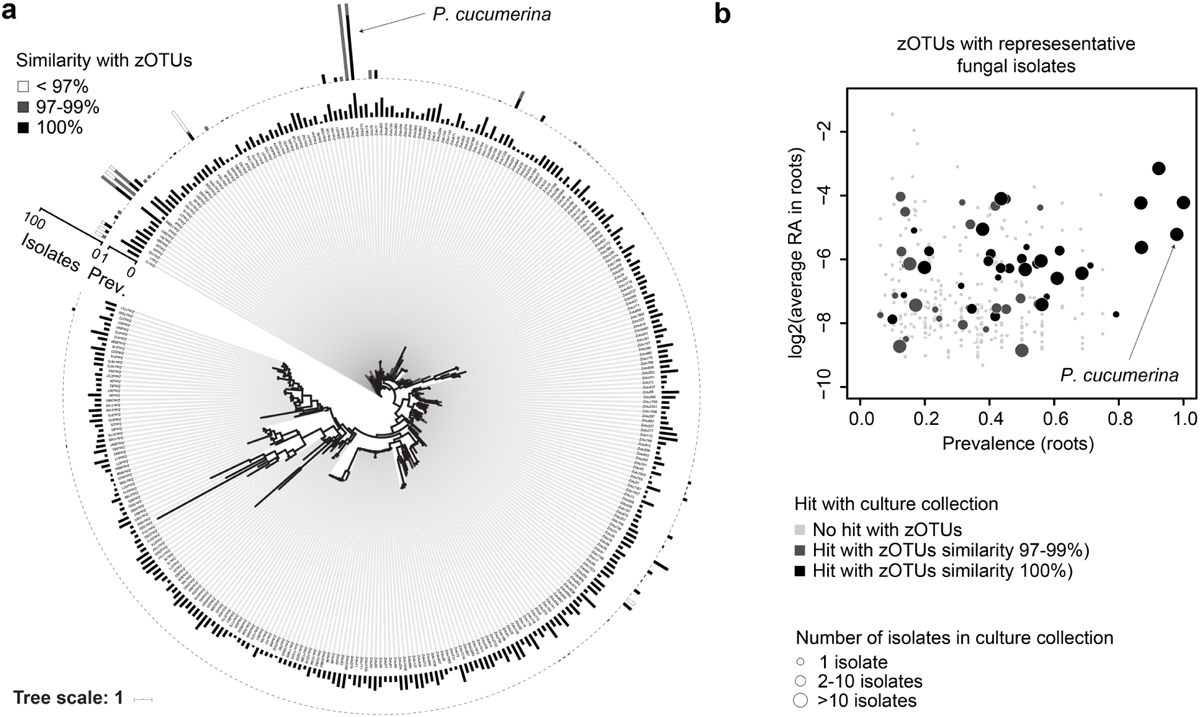
Core fungal taxa detected in Arabidopsis roots across Europe and cross-reference with fungal culture collections. **a,** Maximum likelihood tree based on ITS1 sequences of 338 zero-radius OTUs (Zotus) detected in root samples retrieved from two previous studies (n = 291 samples). This includes *A. thaliana* root samples harvested from 17 European sites and from the Cologne agricultural soil. The inner circle depicts zOTU prevalence across18 sites. The outer barplot depicts Zotus for which representative isolates match culture collection isolates (n = 488 strains, deriving from 4 sites) based on three sequence similarity thresholds (< 97%, 97-99% and 100%). **b,** Zotus are plotted according their mean log2 relative abundance (RA) across all root samples (n = 291 samples), relative to their prevalence across all sample sites (n = 18). RA is only calculated across samples where Zotus were present (i.e., representing at least 0.1% RA per samples). Only root samples with > 1,000 reads were considered for this analysis. Zotus that have hits against the ITS of cultured fungal isolates are colored (dark grey: 97-99% sequence similarity, black: 100% sequence similarity). The size of the circle depicts the number of hits (ranging from 1 isolate to more than 10 isolates).

**Supplementary Fig. 2:**
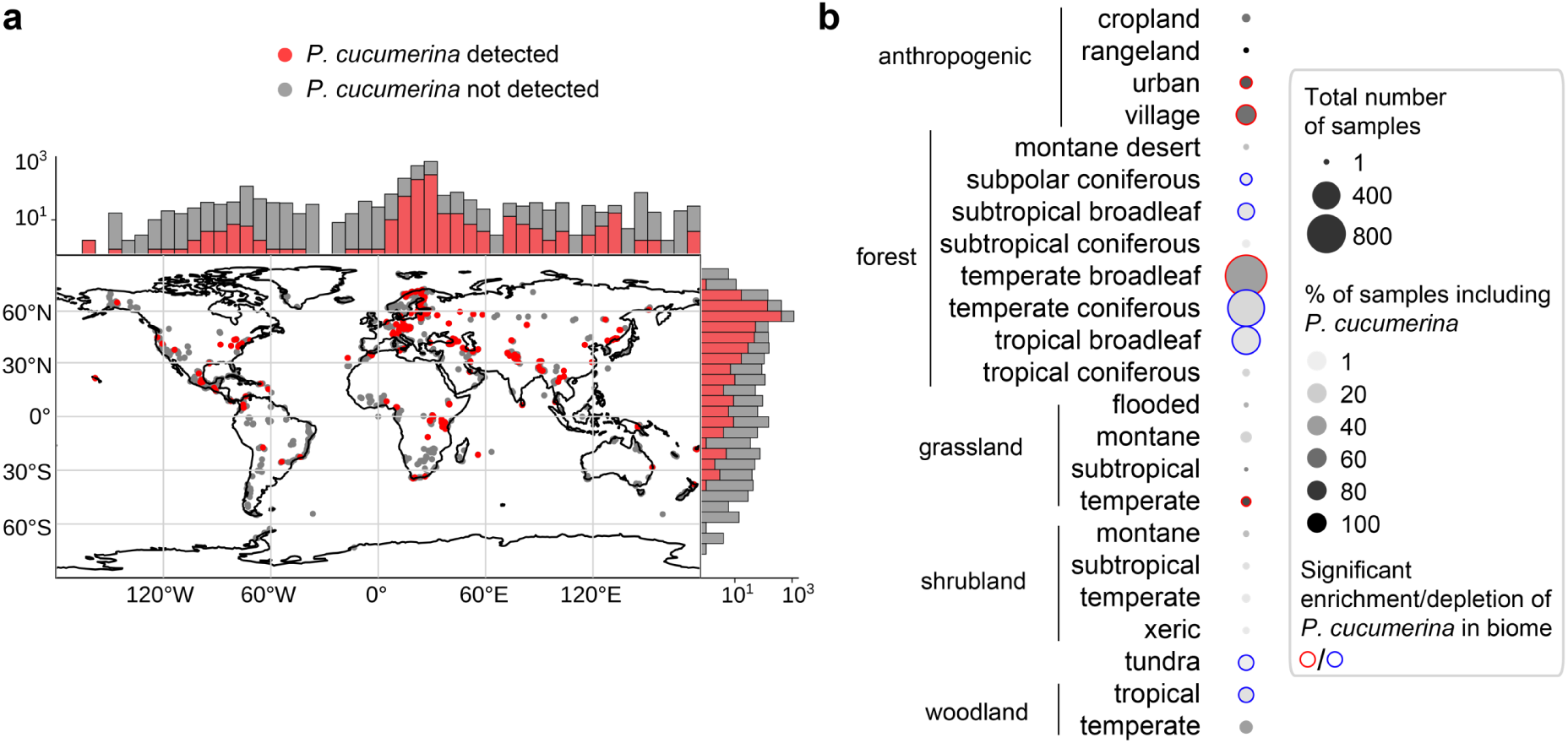
*P. cucumerina* is a ubiquitous soil-borne fungus. **a,** Detection of *P. cucumerina* (presence of curated ITS1 sequences^35^) in soil samples originating from all continents (n = 3,194 samples based on Global Soil Mycobiome dataset^84^). **b,** Soil biome enrichment/depletion in samples containing *P. cucumerina*, as tested by Fisher’s exact test (see Methods) using the Global Soil Mycobiome dataset and the metadata associated to soil samples.

**Supplementary Fig. 3:**
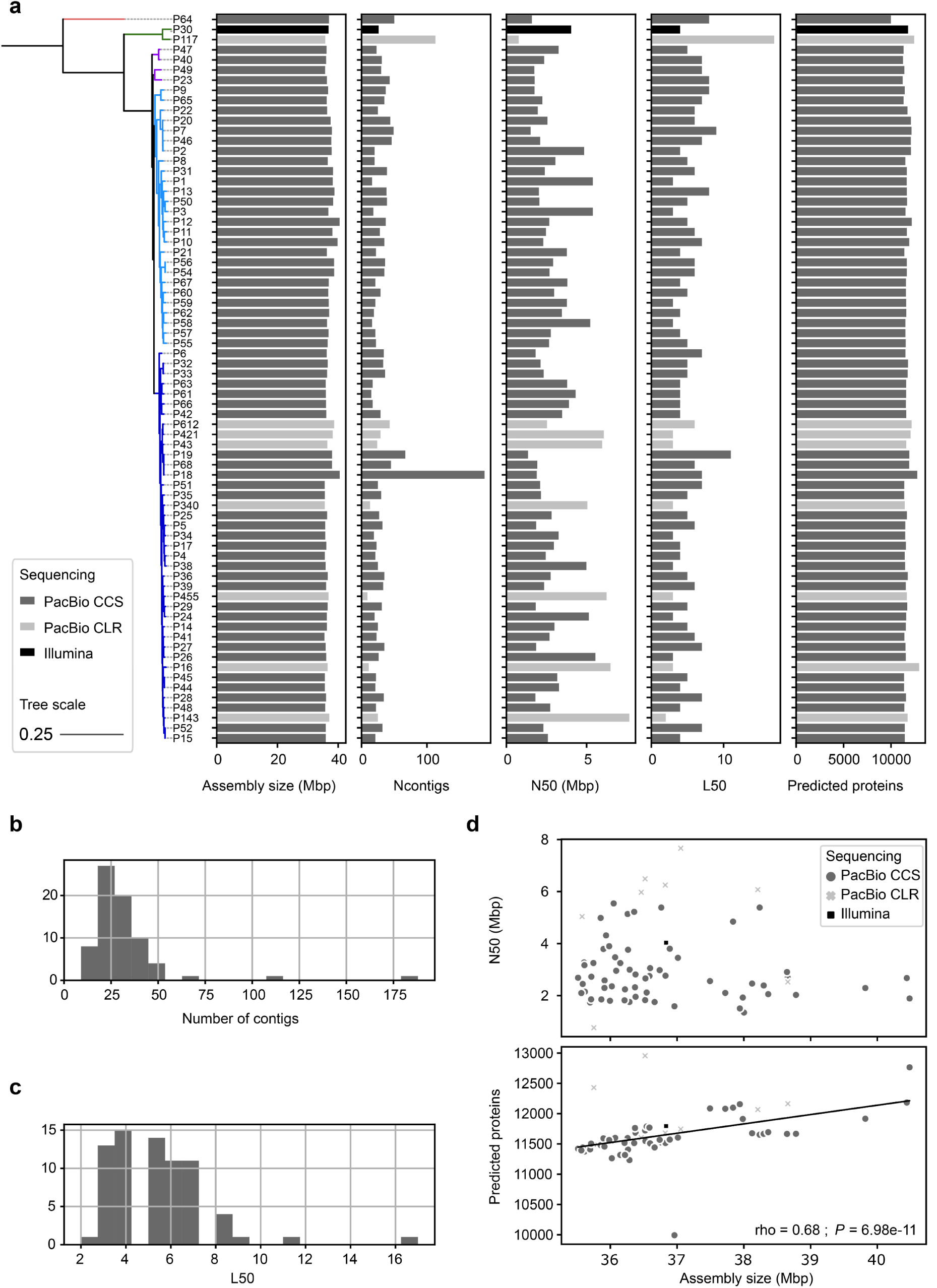
Assembly size and quality for 72 *Plectosphaerella* genomes. **a,** For each genome in our genomic dataset, the following values are plotted: assembly size, number of contigs in the assembly, N50 and L50 values, number of predicted genes in the assembly. These values are represented along the *Plectosphaerella* collection phylogeny reconstructed with IQ-TREE (See Methods). Note that isolates P68 and P16 correspond respectively to *P. cucumerina* BMM (PcBMM) and *P. cucumerina* 0016 (see this work). **b,** Distribution of the number of contigs in the genomic data set (n = 72, one value per genome assembly). **c,** Distribution of the L50 values in the genomic data set (n = 72). **d,** N50 values (top) and numbers of predicted genes (bottom), plotted in function of assembly size. The number of predicted genes is significantly correlated to the assembly size according to Spearman’s rank correlation.

**Supplementary Fig. 4:**
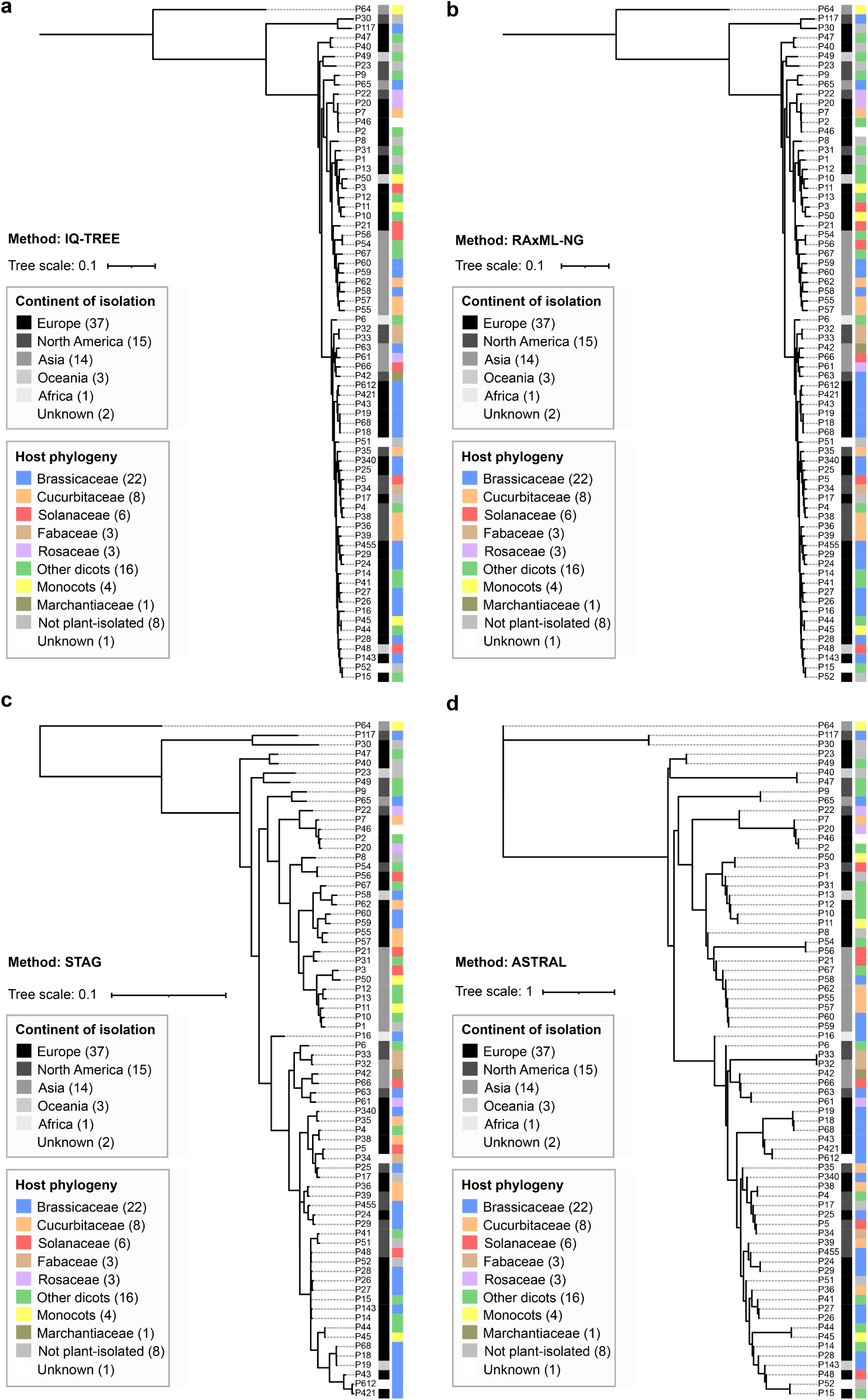
Phylogenomic trees of the 72 *Plectosphaerella* strains, reconstructed with four different methods. **a.** Phylogenetic tree reconstructed with IQ-TREE^100^ (model *JTT+F+I+G4*, as defined by ModelFinder^101^) from the trimmed concatenated alignments of 5,466 single copy orthologs identified by orthology prediction with OrthoFinder^97^. **b.** Phylogenetic tree reconstructed with RAxML-NG^102^ (model *JTT+F+I+G4*, as defined by ModelFinder) from the trimmed concatenated alignments of 5,466 single copy orthologs identified by orthology prediction with OrthoFinder. **c.** Coalescent phylogenetic tree reconstructed from total gene family trees generated by OrthoFinder, with built-in method STAG^133^. **d.** Coalescent phylogenetic tree reconstructed with ASTRAL^104^ from the OrthoFinder-reconstructed gene family trees of 5,466 single copy orthologs. See Methods for further details.

**Supplementary Fig. 5:**
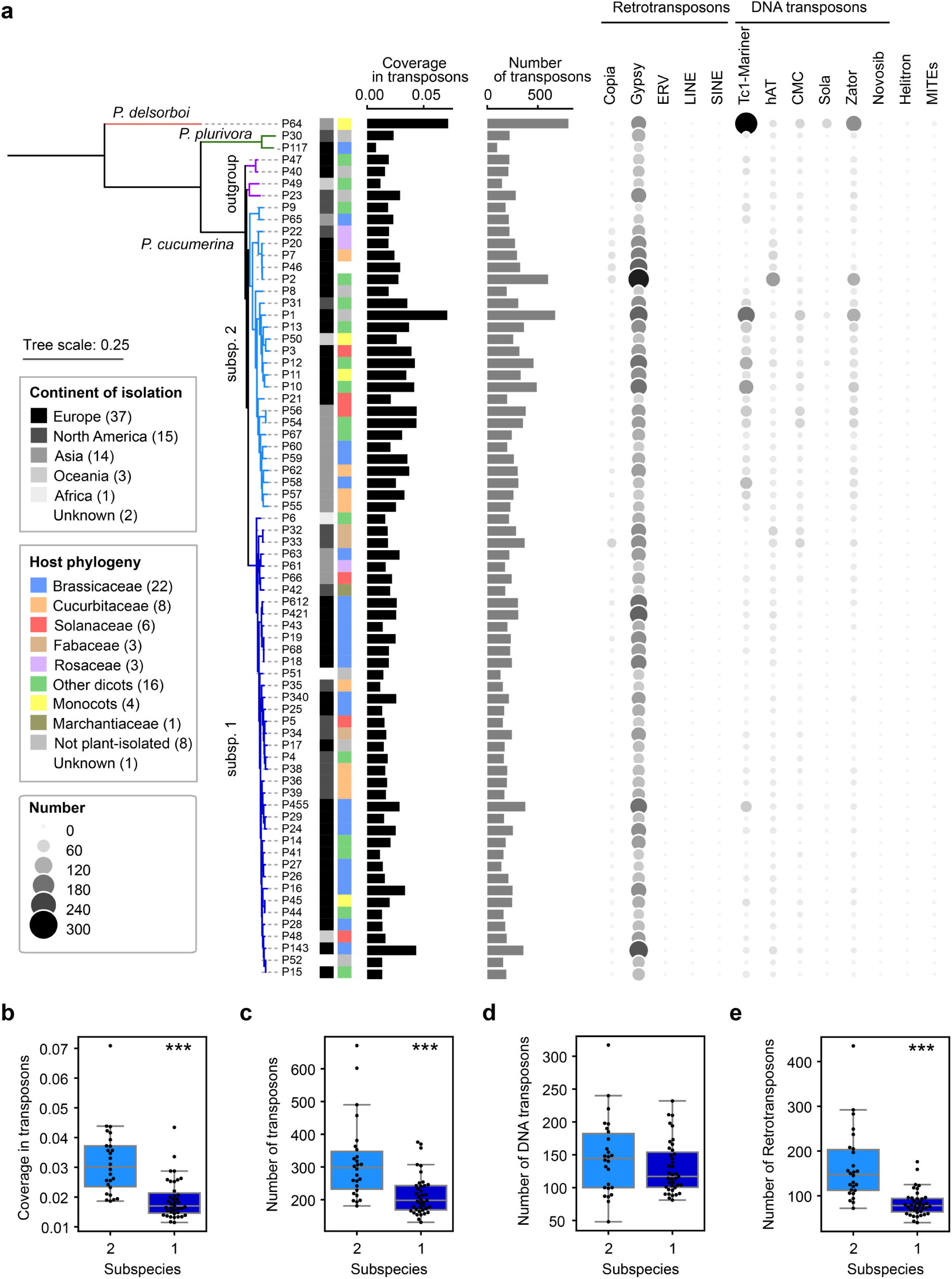
Transposon annotation of the 72 *Plectosphaerella* genome assemblies. **a,** Results of transposon annotation with the pipeline TransposonUltimate^96^. Along the *Plectosphaerella* collection phylogeny, barplots show the percentage of each genome covered by transposons (“coverage”) and the total numbers of transposons annotated. On the right, a bubble plot depicts the number of transposons annotated per family. **b,** Boxplot displaying the difference of *P. cucumerina* genomes in subspecies 1 and 2 in coverage in transposons. Mann-Whitney test revealed a significant difference (statistic = 134.0, *P* = 6.10x10^-7^). **c,** Boxplot displaying the difference of *P. cucumerina* genomes in subspecies 1 and 2 in transposon number. Mann-Whitney test revealed a significant difference (statistic = 194.5, *P* = 2.94x10^-5^). **d,** Boxplot displaying the difference of *P. cucumerina* genomes in subspecies 1 and 2 in DNA transposon number. Mann-Whitney test revealed no significant difference (statistic = 437.0, *P* = 0.35). **e,** Boxplot displaying the difference of *P. cucumerina* genomes in subspecies 1 and 2 in retrotransposon number. Mann-Whitney test revealed no significant difference (statistic = 108.0, *P* = 9.46x10^-8^).

**Supplementary Fig. 6:**
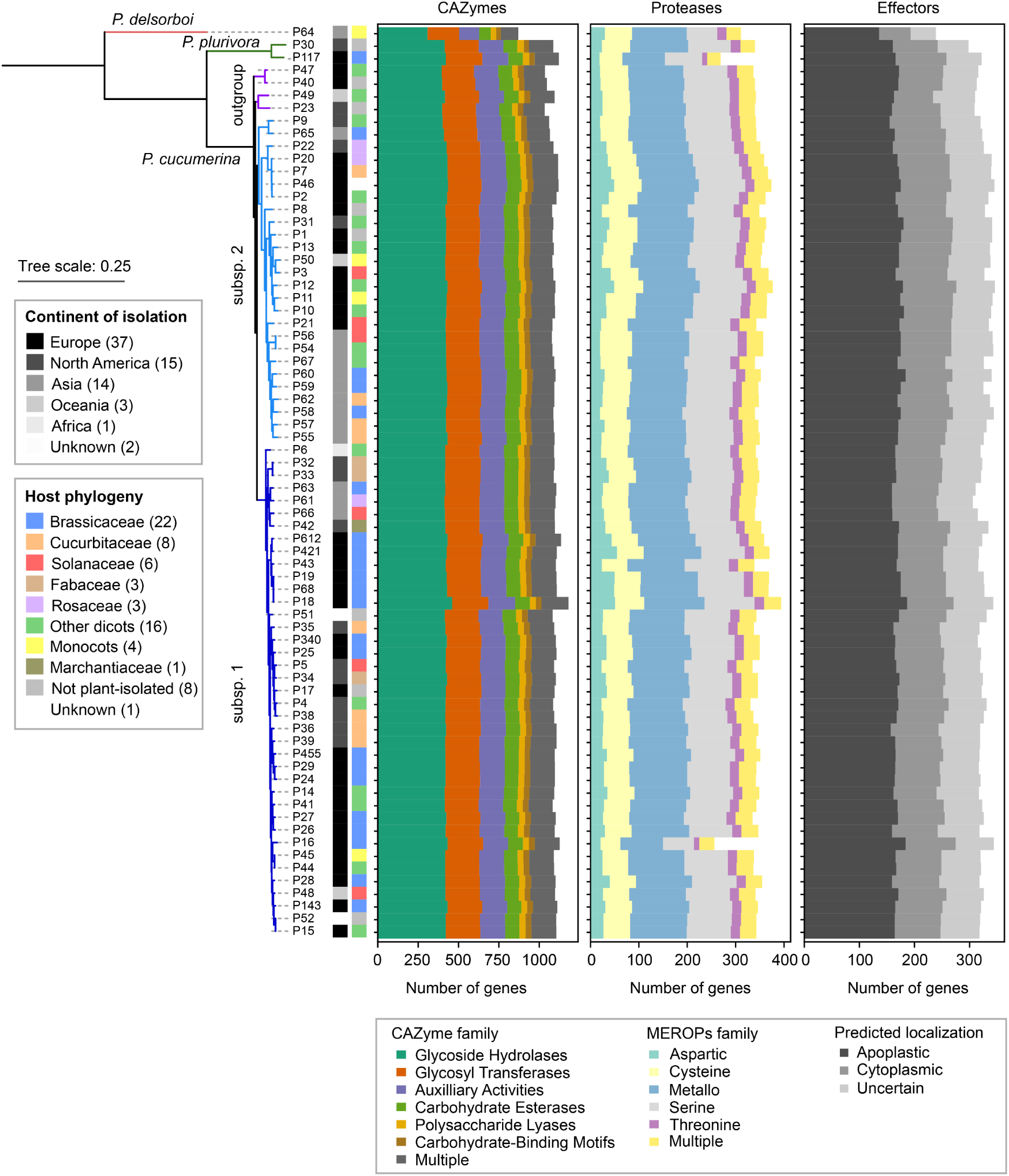
Genome annotation of the 72 *Plectosphaerella* isolates in CAZyme-, protease- and effector-encoding genes. Number of CAZyme-(left), protease-(center) and effector-(right) encoding genes annotated in each of the 72 *Plectosphaerella* genomes, using methods dbcan^47^, emapper^92^ and EffectorP^91^, respectively. Different colors divide the barplots to highlight different functional families.

**Supplementary Fig. 7:**
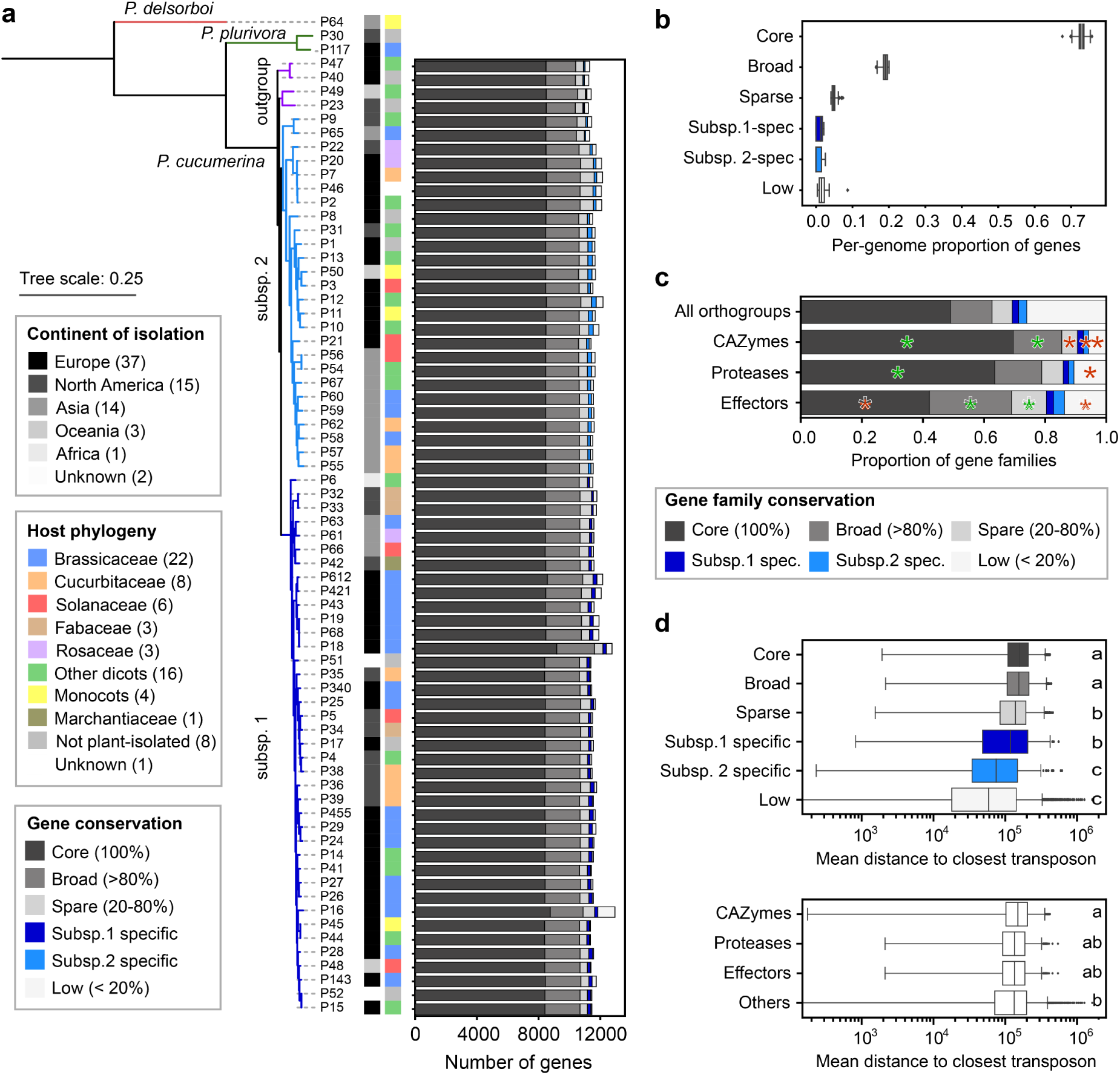
Conservation of gene families across the genomic dataset. **a,** Following orthology prediction (OrthoFinder), gene families (i.e., n = 15,269 orthogroups) were analysed for their conservation across the 69 *P. cucumerina* isolates and classified as core (represented in 100% of *P. cucumerina* isolates), broadly conserved (represented in > 80% of *P. cucumerina* isolates), sparsely conserved (in 20-80% of *P. cucumerina* isolates), subspecies-associated (in one subspecies’ members while absent in the other subspecies) and low-conservation (in < 20% *P. cucumerina* isolates). The number of genes in each category is shown in a stacked barplot, along the *Plectosphaerella* phylogeny. **b,** Per-genome proportions of genes in each one of these categories are shown in a boxplot, revealing that about 75% of *P. cucumerina* gene families are “core” (i.e. conserved across the full collection). **c,** Proportions of total gene families, as well as gene families encoding CAZymes, proteases and effectors, in the different intra-species conservation levels. Fisher’s exact tests comparing proportions of CAZyme, protease and effector gene families to total orthogroups were computed for each conservation level. Significant enrichments/depletions are highlighted with asterisks (green: enrichments, red: depletion). **d,** Mean distance to the closest transposon (in the same contig) were plotted for each gene family, grouped by conservation level (top) or by gene function (bottom). Letters reflect significant differences between groups according to Kruskal-Wallis tests (*P* < 0.05) followed by post-hoc Dunn test.

**Supplementary Fig. 8:**
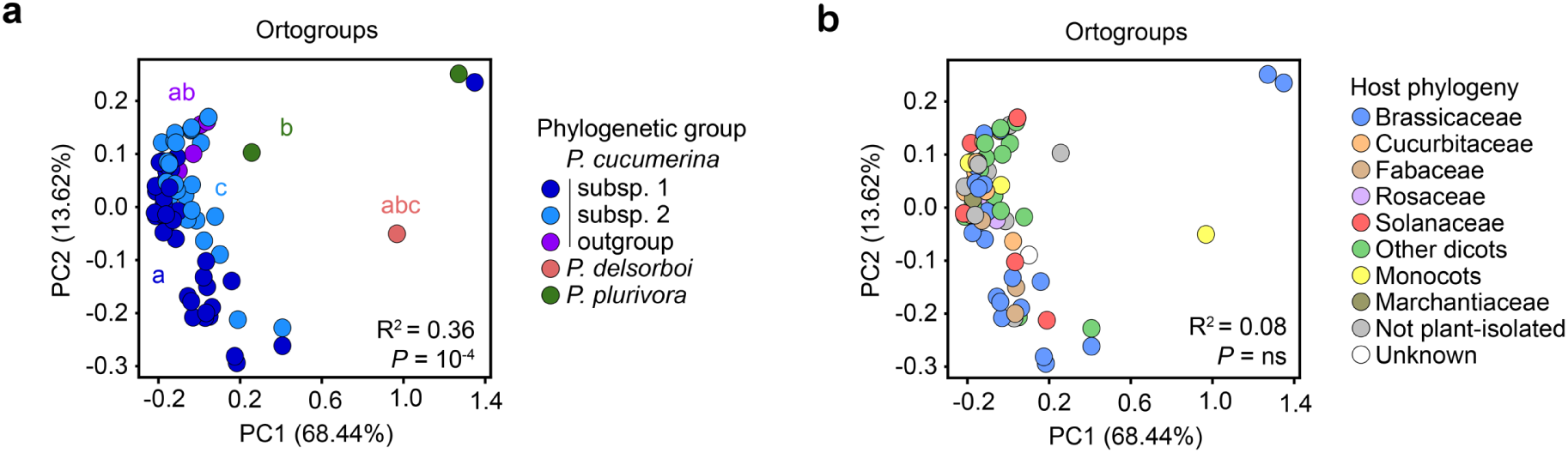
Effect of phylogeny and the host of isolation on inter-strain differences in gene family repertoire. **a,b,** Principal Coordinate Analysis profiling inter-strain differences in total gene family repertoires (= 15,269 orthogroups, OrthoFinder). This PCoA is colored according to strain phylogenetic group (**a**) or host phylogeny (**b**). R^2^ and *P* value resulting from PERMANOVA test assessing the effect of these two categories on the three genomic repertoires are shown at the bottom of each graph.

**Supplementary Fig. 9:**
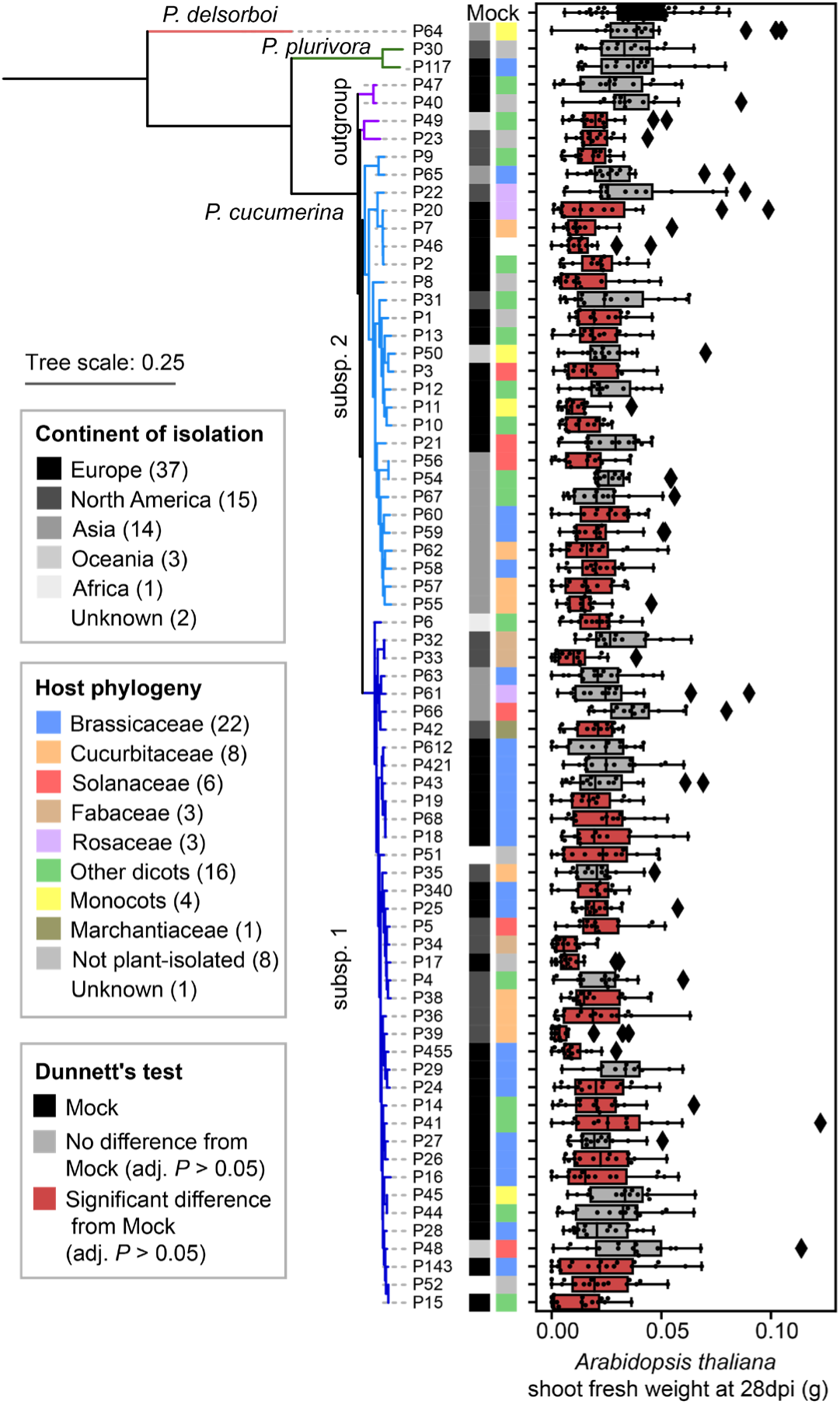
Individual fungal effects of *Plectosphaerella* isolates on *A. thaliana* performance in gnotobiotic system. Shoot fresh weight (SFW) of plants which roots were inoculated with 500 spores of each strain, after 28 days in culture in an agar-based gnotobiotic system (for each fungus: 4 independent biological replicates, 4 plants per replicate). These weight values reflect individual fungal effects on plants. Boxes colored in red reveal a significant difference between the fungus-inoculated and mock-treated plants, according to ANOVA and Dunnett’s test (*P* < 0.05). Boxes are represented along the strain phylogeny, reconstructed with IQ-TREE. Black diamonds represent outlier values. Note that isolates P68 and P16 correspond respectively to *P. cucumerina* BMM (PcBMM) and *P. cucumerina* 0016 (see this work).

**Supplementary Fig. 10:**
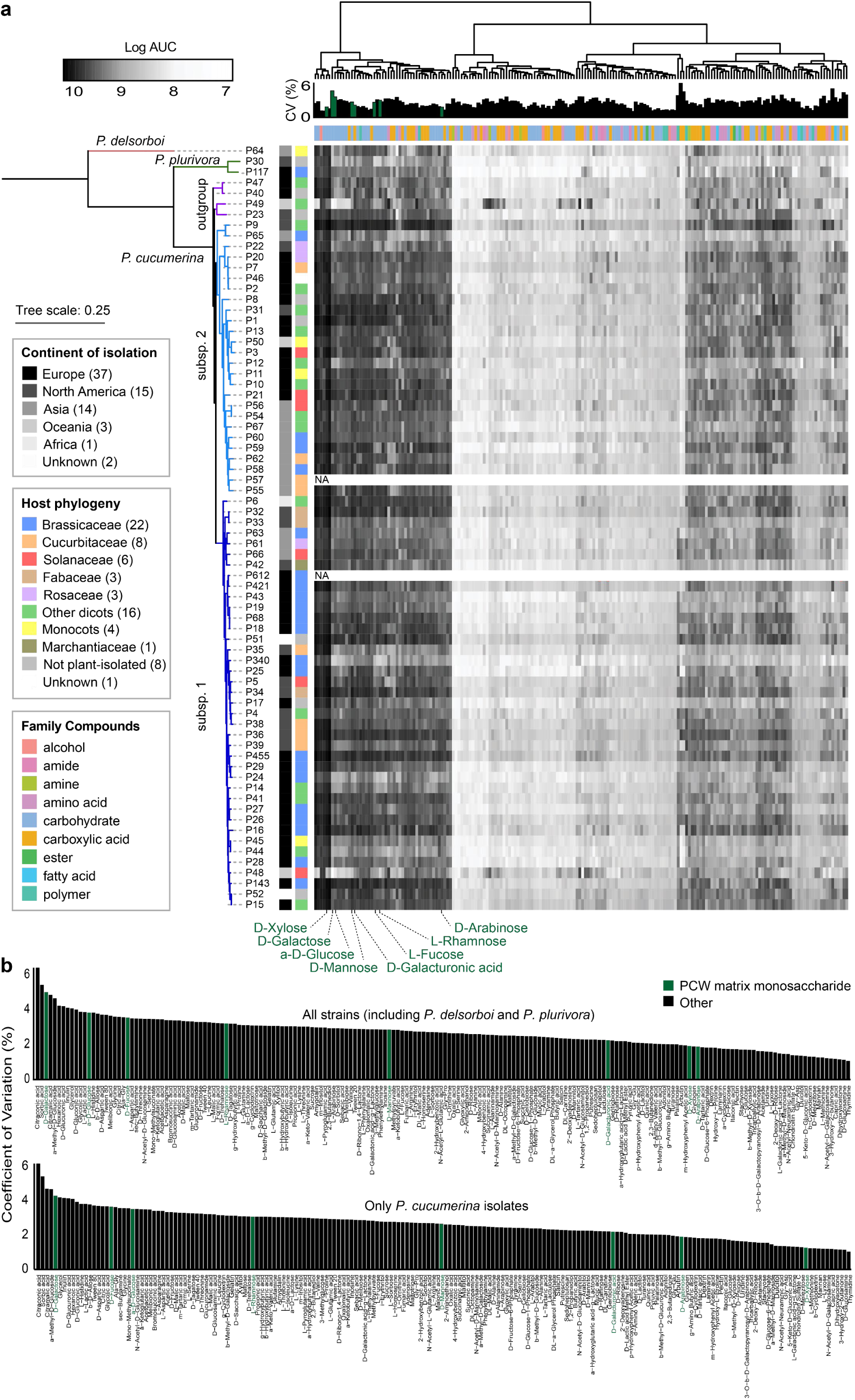
Carbon resource consumption profiles of *Plectosphaerella* isolates. **a,** Heatmap showing the respiration profiles of *Plectosphaerella* strains (n = 72, y-axis) over 5 days when grown on individual carbon sources (x-axis, n = 190) using Biolog PM1 and PM2 plates. The color intensity represents the log-transformed area under the curve (log AUC) of respiration. Carbon sources are clustered based on their utilization patterns across all *Plectosphaerella* strains. Strains are ordered according to their phylogeny. Plant cell wall (PCW) matrix monosaccharides (**Supplementary Fig. 15**) are highlighted in dark green. **b,** Bar plots showing the coefficient of variation (y-axis) in respiration across isolates for each carbon source (x-axis). The top panel includes all strains, while the bottom panel focuses on *P. cucumerina* isolates only. PCW matrix monosaccharides (**Supplementary Fig. 15**) are highlighted in dark green.

**Supplementary Fig. 11:**
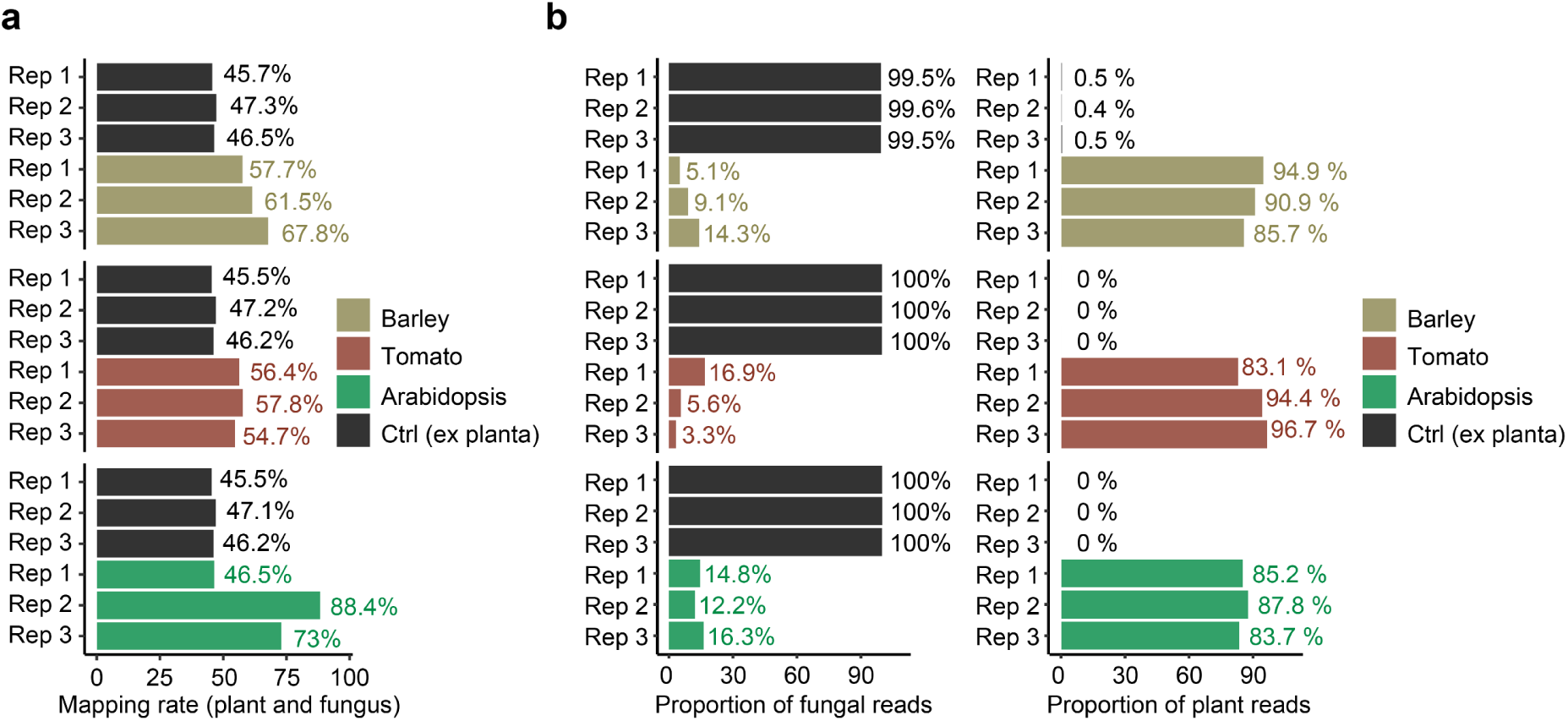
Mapping rates of plant and fungal RNAseq reads against corresponding reference genomes. **a**, The bar chart depicts for each sample the proportion of sequencing reads (i.e., mapping rate) that aligned against the concatenated reference transcriptome of the plant host (either *A. thaliana* Col-0 - TAIR11^109^, Micro-Tom - TMCSv1.2.1^111^, barley - Hvulgare_462_r1^112^ and *P. cucumerina* 0016 utilizing Salmon^113^. **b**, The proportion of fungal reads (left) and plant reads (right) mapped against the corresponding reference transcriptomes. The colors represent the mapping rate of fungal and host reads in roots of barley (khaki), tomato (deep red) and Arabidopsis (green) and of fungal reads in the absence of the host (Ctrl, black).

**Supplementary Fig. 12.**
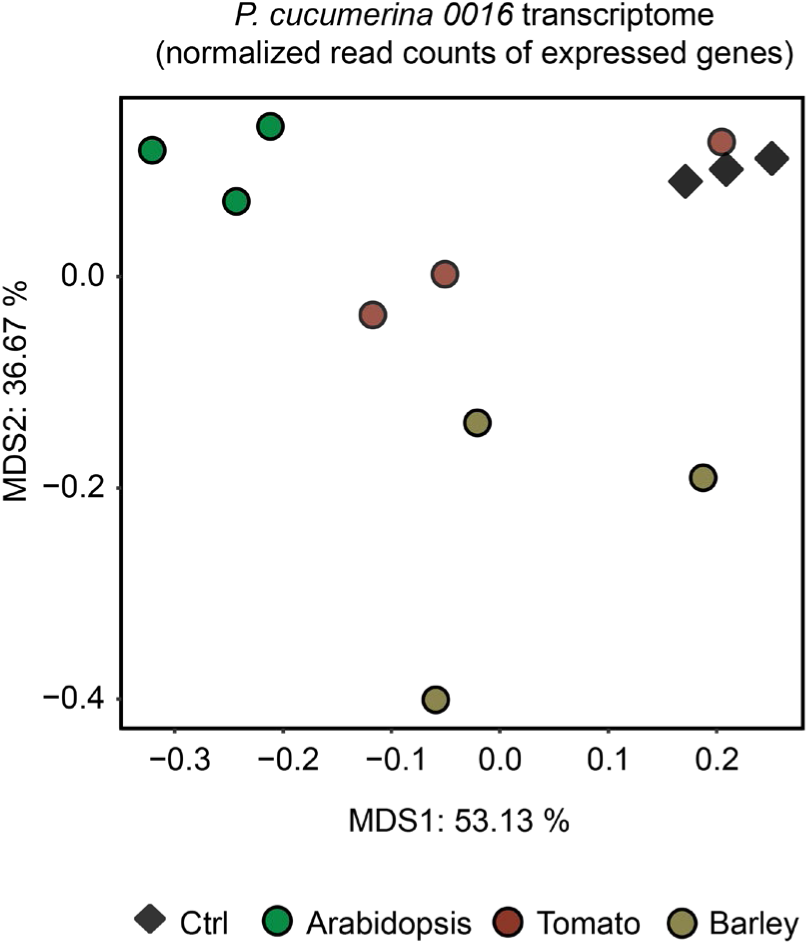
Multidimensional Scaling (MDS) plot of Pearson distances computed on normalized read counts of expressed fungal genes. Fungal transcriptome in roots of Arabidopsis (Green), tomato (Red), and barley (Khaki), as well as in control membrane (Ctrl, *ex planta*, Black). Data points indicate three biological replicates for each condition.

**Supplementary Fig. 13:**
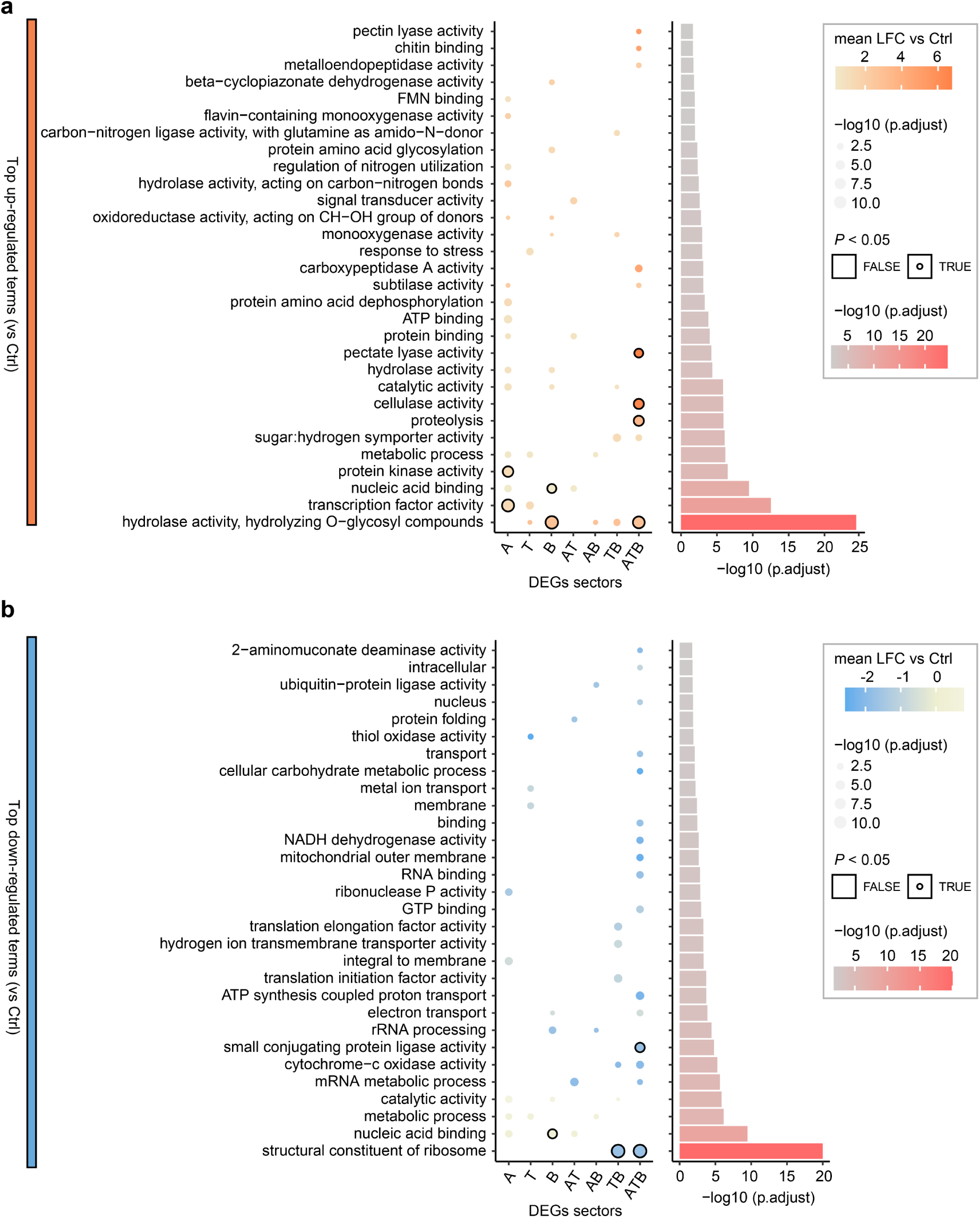
Fungal GO terms significantly regulated during root colonization. **a,b,** The heatmap illustrates the average log_2_ fold-change values for the categories (x-axis), marked circles signify *P* ≤ 0.05 and the size of the circle indicates -log_10_ *P*-adjusted values (FDR, Fisher’s Exact Test). The corresponding barplot shows the cumulative -log_10_ *adj. P* values (x-axis and colors grey-red) for the 30 most common GO-terms in y-axes. GO terms that are up-regulated are represented in orange (**a**) and those that are down-regulated are represented in blue (**b**).

**Supplementary Fig. 14:**
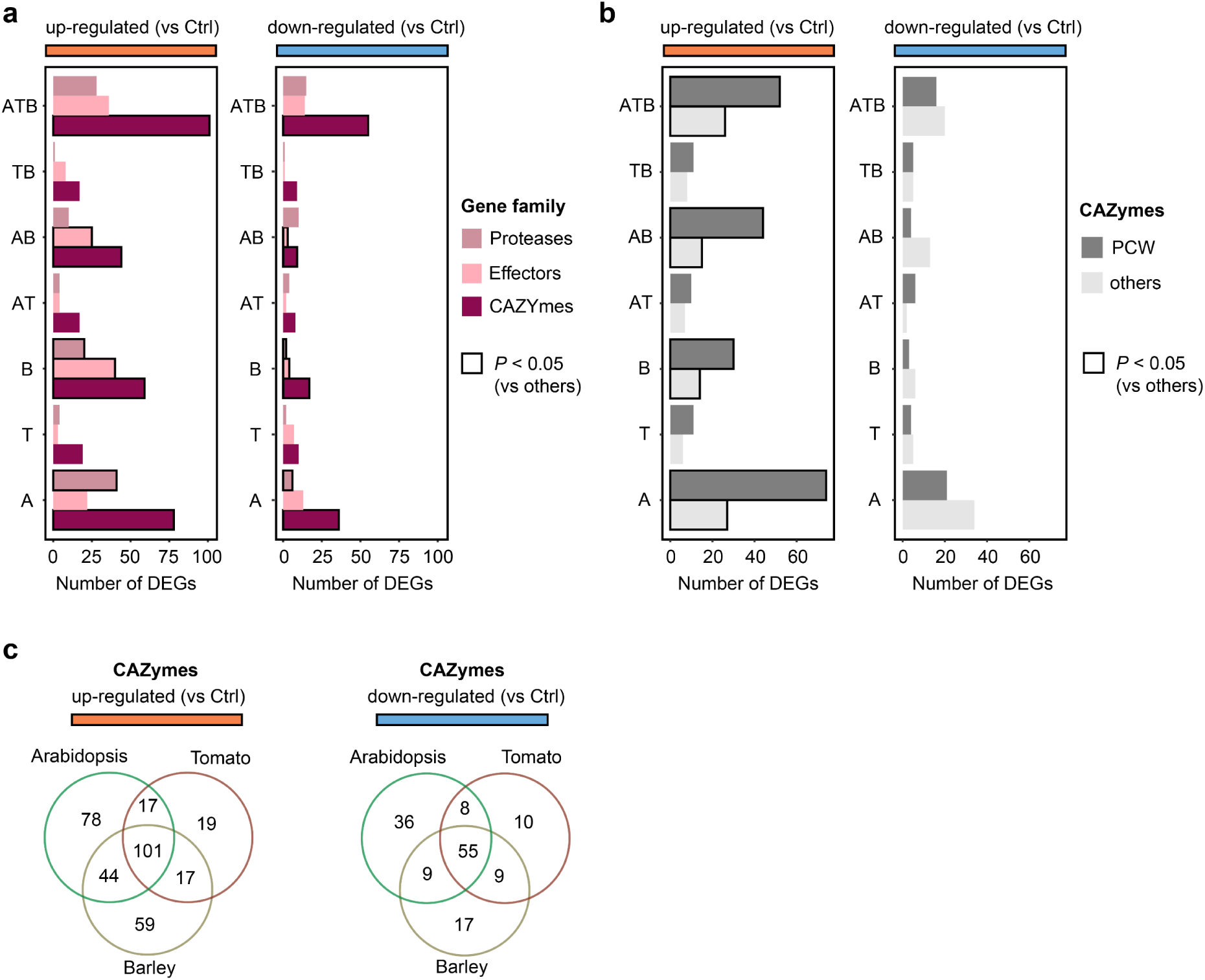
Regulation of CAZyme-encoding genes during root colonization. **a,b,** The combination of barplot illustrates the number of differentially expressed genes (DEGs, x-axes) that are either upregulated (left, denoted in orange) or downregulated (right, denoted in blue) for the shared sectors on the y-axes. Marked bars signify those sectors that are *P* ≤ 0.05 (prop.test, chi-square statistics) indicating the ratio of gene family to be elevated among the total DEGs in the dataset. The colors of the barplot elucidate the predetermined gene families Proteases (deeppink), Effectors (lightpink) and CAZYmes (magenta) (**a**) and in grey plant-cell-wall components (PCW; mined “pectin, cellulose, lignin, arabinan, galactan, glycan, glucan, xylan, galacturon) CAZY-Substrate (dbCAN^47^) (**b**). **c**, Venn diagram illustrating the number of DEGs encoding CAZymes (|log2FC| ≥ 1 and FDR ≤ 0.05) that are up-regulated (orange) and down-regulated (blue) *in planta*, respectively.

**Supplementary Fig. 15:**
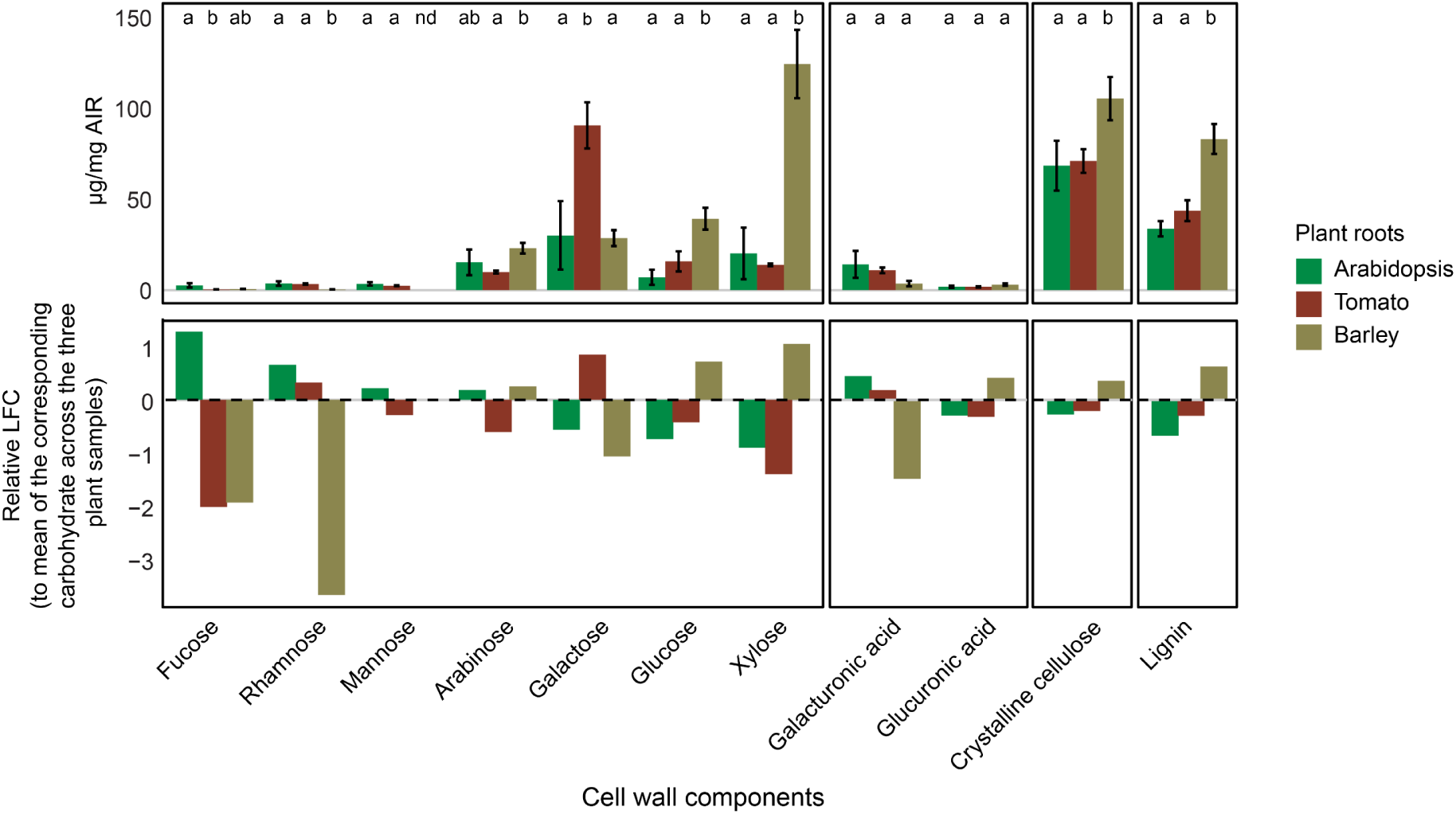
Root cell wall composition analysis of Arabidopsis, tomato, and barley: The upper bar plot illustrates the concentration of wall matrix monosaccharides, including uronic acids [in µg/mg alcohol insoluble residue AIR, y axis], cellulose, and lignin detected in roots of 5-week-old *A. thaliana* (n = 36 plants) and tomato (n = 18 plants), and 4-week-old barley (n = 6 plants). The cell wall components were arranged based on the different assays used and in descending order based on their abundance in Arabidopsis. Multiple pairwise comparisons between the three plant species for each cell wall component were performed using Kruskal-Wallis-Dunn Test, adjusted with Bonferroni correction (*P* ≤ 0.05). The lower bar plot illustrates the relative enrichment of carbohydrates (log2FC) for each plant with respect to the mean value measured across all three plants. Data derived from three biological replicates.

**Supplementary Fig. 16:**
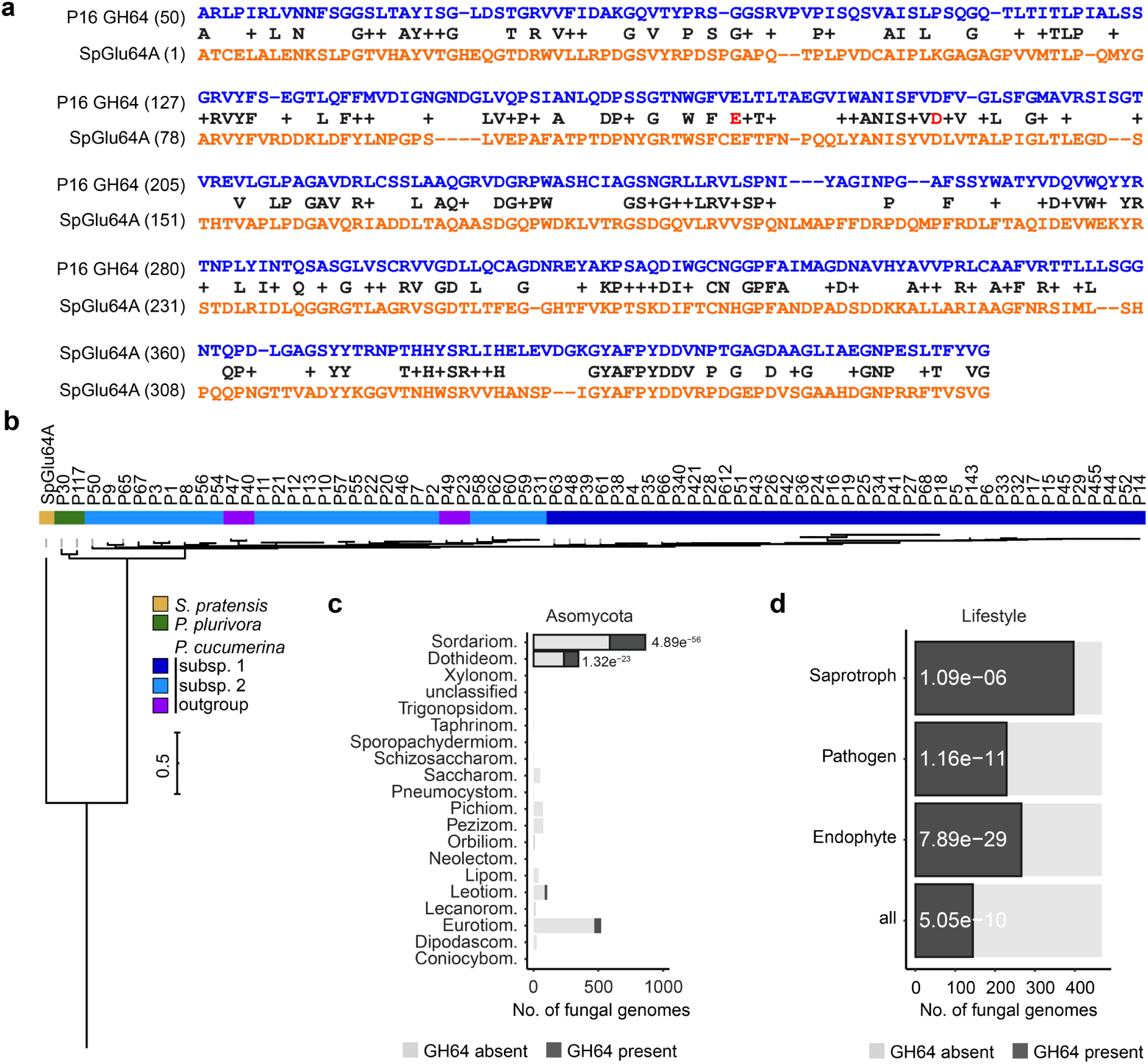
Function and evolutionary conservation of GH64 in the fungal kingdom. **a,** Structure-aware sequence alignment of *P. cucumerina* 0016 GH64 (in blue) with SpGlu64A (in orange) shows conservation of catalytic residues (in red), despite overall low sequence similarity (sequence identity=34.3%). **b,** Maximum-likelihood phylogenetic tree (model JTT+R2 of iqtree^100^) of the orthologs of *P. cucumerina* 0016 GH64 in the *Plectosphaerella* genome collection, together with SpGlu64A. Labels refer to the strain encoding the protein, while color strips highlight the strain phylogenetic group. Note: except for *P. delsorboi* which does not encode any GH64 enzyme, a single GH64-encoding gene was annotated in every *Plectosphaerella* genome. **c,d,** Stacked bar plot depicting the presence/absence of GH64-encoding genes in 2,534 fungal genomes (x-axes) across distinct classes of Ascomycota members (**c**) and fungal lifestyles (**d**) (y-axes). Highlighted in dark grey are those where *GH64* family genes are encoded in the genome, and those with thick black borders are enriched within the group (Fisher’s exact test followed by *P* value adjustment using Bonferroni correction).

**Supplementary Fig. 17:**
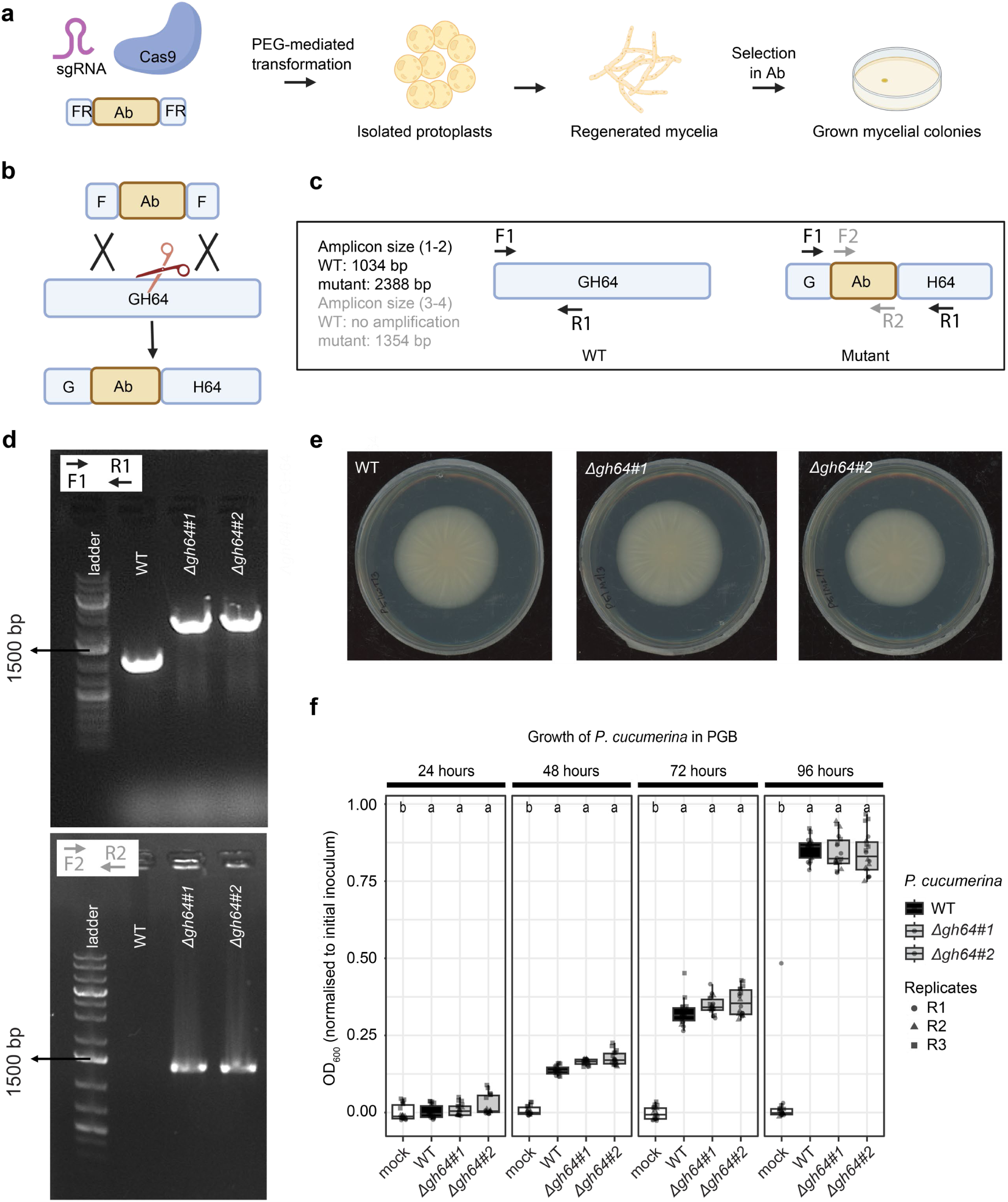
Protoplasts, regenerated Protoplasts and *P. cucumerina* 0016 mutants validation: **a**, Schematic diagram illustrating PEG-mediated transformation and CRISPR/Cas9-mediated knockout method. (PEG-mediated transformation method: 1. Isolation of protoplasts, 2. transformation of Donor DNA and Cas9-sgRNA complex into the protoplasts. 3. Regeneration of protoplasts. 4. Selection of potential colonies in media containing Phleomycin.) **b,** The CRISPR/Cas9 system enhances HR-mediated integration by introducing a double-stranded break in the target DNA, allowing for effective integration with short homology flanks of 60bp. **c**, Schematic representation of the *gh64* gene region in WT and in mutant fungi. Primer pairs used in panel d are depicted. **d**, Targeted disruption of the GH64 gene. Top: Agarose gel electrophoresis results using primer pair F1, R1. The wild type shows a band at 1034 bp, while mutants display a band shift to 2388 bp due to the insertion of the antibiotic resistance sequence. Lane 1: Ladder; Lane 2: Wild type (WT) control; Lane 3, 4: Mutant 1, 2; Bottom: Amplification of the phleomycin antibiotic resistance cassette using primer pair F2, R2. The gel schema is identical to the top panel, confirming the presence of the phleomycin cassette in the mutant lines. **e,** *In vitro* growth of *P. cucumerina* 0016 WT *versus Δgh64* mutant strains in PGA plate. **f**, Dynamics of *in vitro* growth of *P. cucumerina* 0016 WT *versus Δgh64* mutant strains in PGB medium.

**Supplementary Fig. 18:**
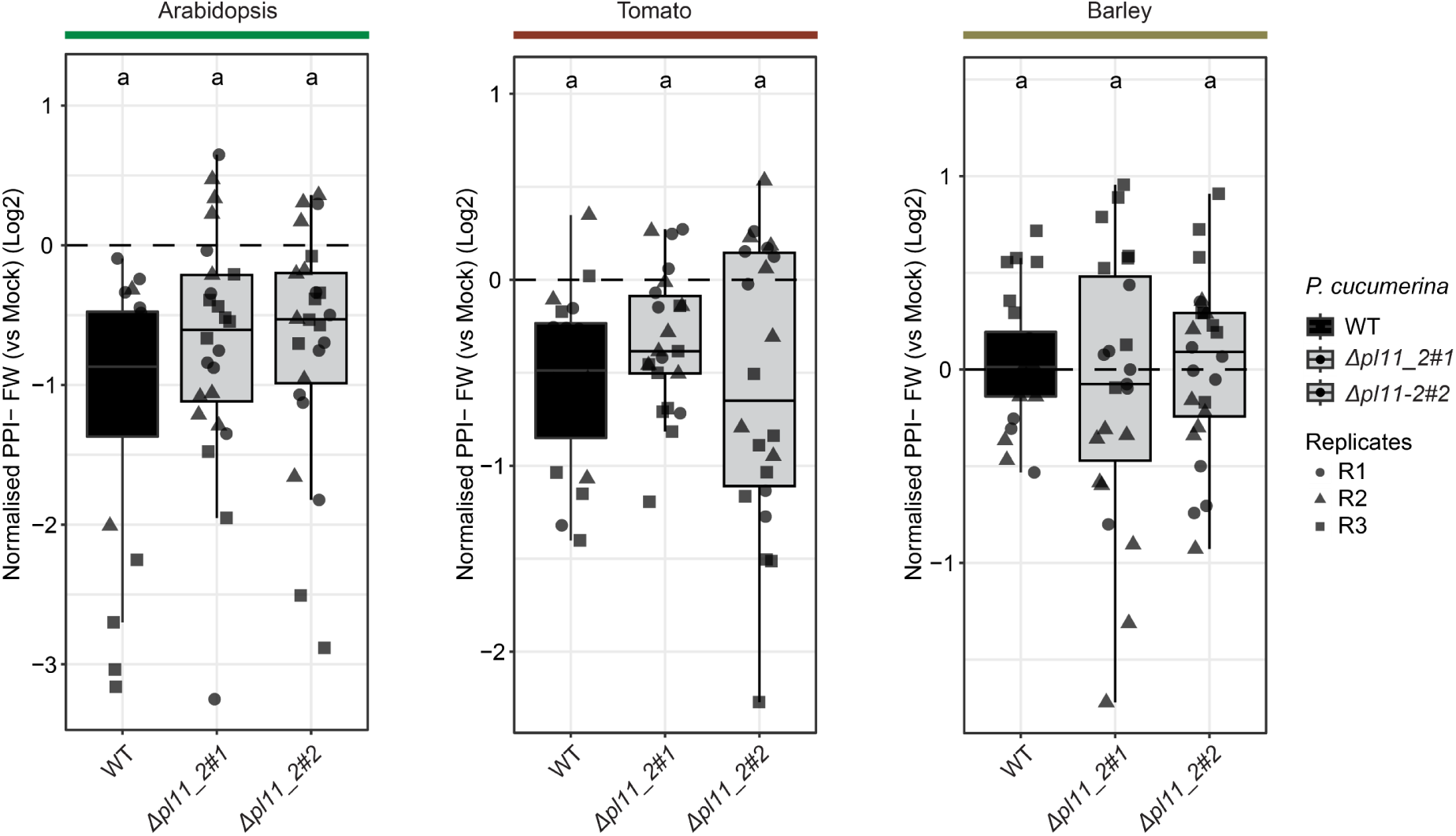
Phenotype assessment of P. cucumerina 0016 Δpl11_2 mutant strains. Plant Performance Indices (PPI) of Arabidopsis, tomato, and barley colonized by *P. cucumerina* 0016 WT and mutant strains (*Δpl11_2#1*, *Δpl11_2#2*) were normalized to mock-treated plants and log2 transformed. PPI values were calculated by multiplying the germination rate by the shoot fresh weight of the plants. Pre-grown seedlings (2-week-old Arabidopsis and tomato and a week-old barley) were inoculated with *P. cucumerina* 0016 and harvested 21 days post fungal inoculation. Three independent biological replicates are depicted (circles, triangles, squares), each consisting of 5-8 plants (n = 15-24 plants per condition). Multiple pairwise comparisons of plant performance between mock-treated and *P. cucumerina* 0016 treated plants were performed using Kruskal-Wallis-Dunn Test, adj. with Bonferroni correction (*P* ≤ 0.05).

**Supplementary Fig. 19:**
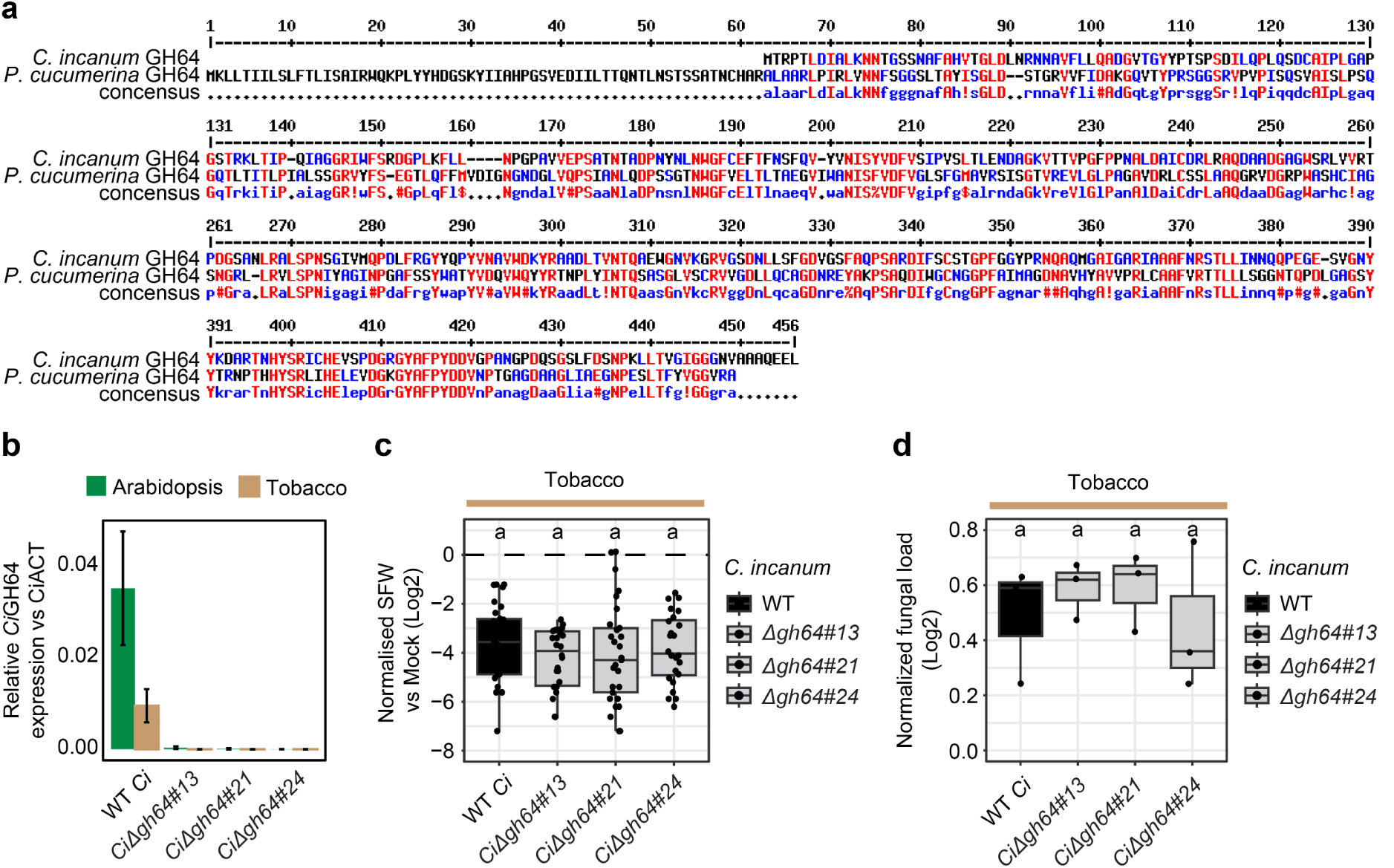
*C. incanum* GH64 sequence conservation, expression and functional relevance for tobacco root infection. **a,** Protein sequence alignment of *P. cucumerina* 0016 GH64 (bottom) with *C. incanum* GH64 (up) obtained from Multalin^134^. Note the absence of signal peptide in *C. incanum* GH64 sequence. **b**, *GH64* transcript quantification in roots of Tobacco and Arabidopsis. The relative expression of the *C. incanum GH64* gene was measured by qRT-PCR on tobacco or Arabidopsis roots inoculated with *C. incanum* WT (WT *Ci*) and *Δgh64* mutant lines and normalized to the expression of *C. incanum ACTIN* gene at 24 dpi. Statistical significance was evaluated by Tukey’s post-hoc test indicated by different letters, *P* < 0.05. **c**, Shoot fresh weight (SFW) of tobacco plants inoculated with WT *Ci* or *Δgh64* mutants and normalized to mock-treated plants (24 dpi). Statistical significance was evaluated by Tukey’s post-hoc test indicated by different letters (*P* < 0.05). **d**, Relative fungal load quantification of WT and mutant lines in tobacco roots was estimated by qRT-PCR based quantification of *C. incanum ACTIN* transcripts normalized to tobacco *ACTIN* transcripts (24 dpi). Statistical significance was evaluated by Tukey’s post-hoc test indicated by different letters, *P* < 0.05.

## Notes

### Competing Interest Statement

The authors have declared no competing interest.

### Summary of Updates

- One author is removed from the list. At the beginning, the co-authorship was accepted. However, the policy requirements has changed at their institute, so it was not possible to be a part of the publication anymore. - Funding number of A.K.B is updated.

